# Neural substrates of evidence accumulation in social-affective decision-making under perceptual ambiguity

**DOI:** 10.1101/2025.06.24.660932

**Authors:** Sai Sun, Tao Xie, Yibei Chen, Jinge Wang, Xin Li, Rongjun Yu, Peter Brunner, Jon T. Willie, Shuo Wang, Hongbo Yu

## Abstract

Evidence accumulation models have been widely used to explain decision-making in sensory and cognitive domains, but how this process explains social-affective decision-making under perceptual ambiguity remains unclear. Here, we integrate computational modeling with multimodal neuroscience to investigate how perceptual ambiguity in emotion judgment influences decision dynamics. Participants viewed ambiguous stimuli—morphed images blending emotional expressions—and made binary categorizations. Using drift diffusion modeling (DDM), we show that drift rate, a core index of evidence accumulation, decreases as ambiguity increases. Scalp EEG data reveal that the late positive potential (LPP) tracks drift rate in both emotional and non-emotional decisions, but only when perceptual ambiguity is relevant to the task. Likewise, single-unite recordings from the dorsomedial prefrontal cortex (dmPFC) and amygdala neurons show that trial-by-trial firing rates in both regions encode drift rate, with opposite encoding patterns. fMRI-based connectivity analyses further reveal that the strength of amygdala–dmPFC coupling correlates with individual differences in the modulator effect of perceptual ambiguity on drift rate. Together, these findings identify a corticolimbic network that dynamically modulates evidence accumulation during social-affective decision-making under perceptual ambiguity. By linking computational parameters to neural activity across multiple modalities, this work advances our understanding of how the brain resolves emotionally ambiguous perceptual information. It provides a translational framework for studying emotion recognition and decision-making impairments in clinical populations.

## Introduction

Faces are central to social communication, conveying emotionally and socially salient information essential for successful interaction. A core process in interpreting facial expressions is categorical judgment, which likely relies on evidence accumulation, a dynamic mechanism wherein information is integrated over time until a decision threshold is reached. Evidence accumulation is well characterized in sensory domains such as odor, color, and motion perception (Brosnan et al., 2020; Gherman et al., 2024; Hanks & Summerfield, 2017; Kelly & O’Connell, 2013; O’Connell & Kelly, 2021; S. P. Kelly et al., 2021; Tavares et al., 2017; Zhou & Freedman, 2019), as well as in value-based economic and social decision-making (H. Yu et al., 2021; Hutcherson et al., 2015; M. A. Pisauro et al., 2017; M. Milosavljevic et al., 2010; Polanía et al., 2014). In facial emotion perception, however, decision evidence often reflects observers’ interpretations of ambiguous emotional cues based on internal criteria rather than objective quantities such as motion direction, color, or monetary value.

Drift diffusion modeling (DDM) provides an established framework for modeling such decision dynamics (Forstmann et al., 2016; H. Kim et al., 2026, 2026; Mulder et al., 2014; O’Connell et al., 2018; O’Connell & Kelly, 2021; Ratcliff & Smith, 2004; Roberts & Hutcherson, 2019). Previous work has linked DDM parameters to sensorimotor, prefrontal, and parietal systems using EEG, fMRI, and invasive recordings across species (Brosnan et al., 2020; Gherman et al., 2024; Hanks & Summerfield, 2017; Noppeney et al., 2010; Rahnev et al., 2016; Scott et al., 2017; V. De Lafuente et al., 2015). Importantly, DDM has also been applied to affective perception and decision-making (Roberts & Hutcherson, 2019)(Roberts & Hutcherson, 2019). For example, Yau et al. (2020) showed that fusiform representations of facial emotion modulate drift rate during dynamic happy–sad categorization (Yau et al., 2020), and Yau et al. (2021) showed that centroparietal positivity tracks evidence- and urgency-related signals in a similar face-morphing task (Yau et al., 2021). Haller et al. (2024) used DDM and fMRI to distinguish sensitivity and bias during happy–angry face-emotion labeling (Haller et al., 2024), while El Zein et al. (2024) showed prioritized neural processing of socially threatening cues during perceptual decision-making (Zein et al., 2024). Thus, DDM is already an important tool for studying social-affective decisions.

Building on this literature, the present study uses DDM as a common computational framework for integrating neural measures previously implicated in ambiguity processing. Prior work using overlapping datasets showed that the late positive potential (LPP) tracks perceptual ambiguity in facial emotion judgment (Sun, Yu, et al., 2017a; Sun, Zhen, et al., 2017), that human amygdala neurons parametrically encode emotional intensity and categorical ambiguity (S. Wang et al., 2017), and that amygdala–medial frontal connectivity is associated with ambiguity processing (Sun, Yu, et al., 2023). What remains less clear is whether these ambiguity-related neural signals map onto a shared latent decision process—namely, evidence accumulation efficiency. This computational framework is useful because accuracy and reaction time alone cannot distinguish slower evidence accumulation from changes in decision threshold. DDM allows these mechanisms to be compared by estimating latent parameters such as drift rate and boundary separation. Thus, our central question is whether ambiguity-related behavioral and neural effects are best explained by changes in evidence accumulation.

Here, we applied DDM to choice and reaction time data from categorization tasks involving emotionally ambiguous facial expressions and control stimuli. We first tested whether perceptual ambiguity is best captured by reduced drift rate rather than altered boundaries, and whether this effect depends on task relevance. We then examined whether LPP amplitude, single-neuron activity in the amygdala and medial frontal cortex, and amygdala–medial frontal functional connectivity are associated with DDM-derived evidence accumulation. By integrating behavioral modeling with EEG, single-unit recordings, and fMRI, we provide a computational synthesis of how neural signals related to social-affective ambiguity map onto latent decision dynamics, while remaining cautious about the correlational nature of the neural evidence.

## Methods

### Overview of Experiments

In our main task, participants viewed face images morphed along a continuum between happy and fearful expressions (**Figure 1a**). Each stimulus fell into one of seven graded levels: 100%/0% happy (unambiguity), 70%/30% happy (medium ambiguity), 60%/40% and 50% happy (high ambiguity). Participants were asked to categorize each face as either happy or fearful by pressing the corresponding button. A previous study using the identical task has shown that as the ambiguity level increases, participants’ responses become nosier and slower (For the basic behavioral and ERP results, please see (Sun, Zhen, et al., 2017)).

**Figure 1.**
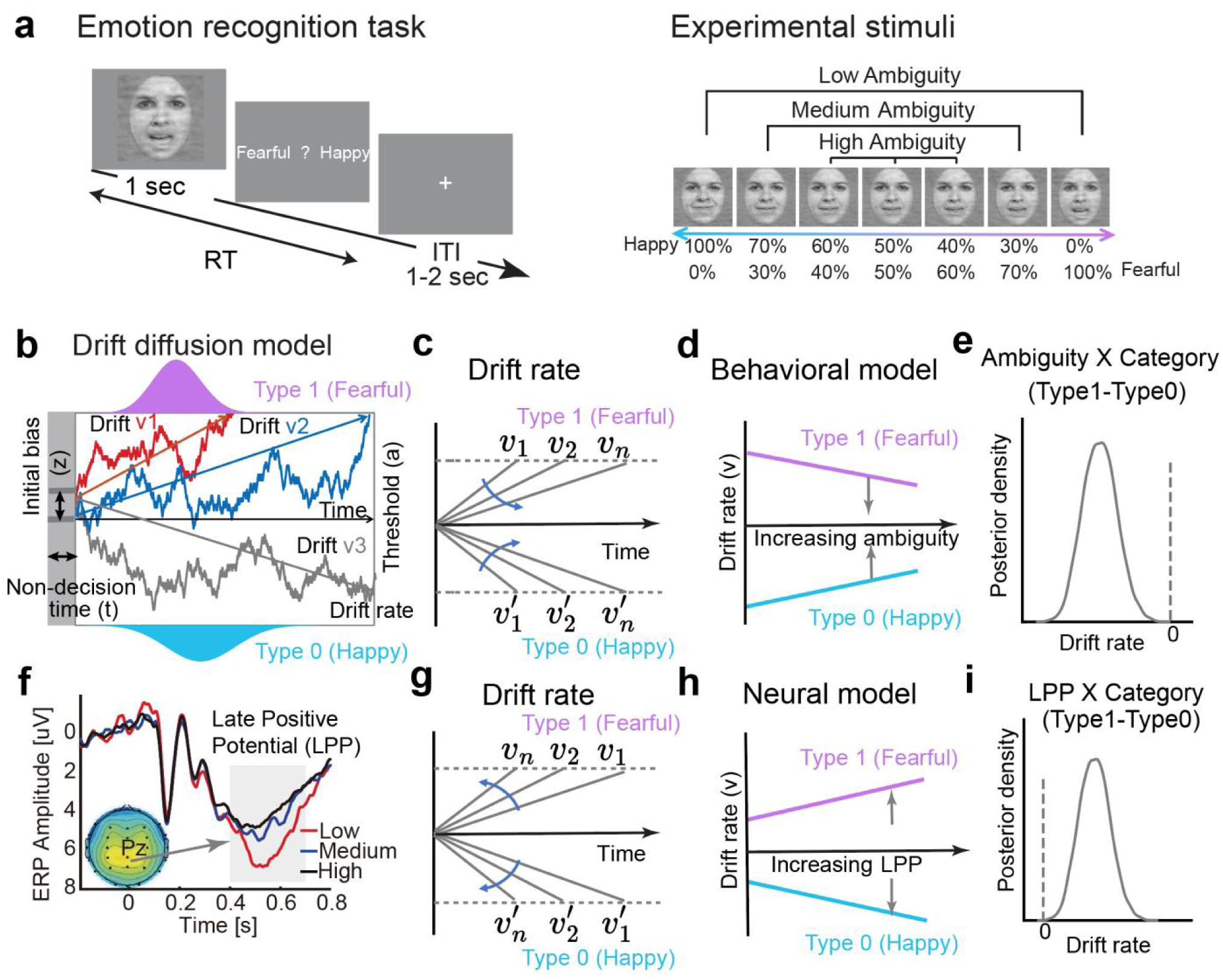
Overview of tasks, DDM models, and hypotheses. **(a)** Perceptual stimuli and task procedure. Participants viewed facial expressions that were morphed along a continuum between fearful and happy facial expressions. The stimuli consisted of seven graded levels, with low ambiguity at the extremes (100% happy or fearful), moderate ambiguity at intermediate morphs (e.g., 70% happy or fearful), and high ambiguity at the midpoint (50%) and near-midpoint levels (40% and 60% happy or fearful). Participants were asked to categorize each face as either happy or fearful by pressing the corresponding buttons. Note that the facial images shown here are adopted from a publicly available database ^63^. **(b)** DDM characterizes decision-making as a noisy evidence accumulation process toward one of two decision boundaries: the upper boundary (Type 1) corresponds to “happy” responses, while the lower boundary (Type 0) corresponds to “fearful” responses. Simulated trajectories illustrate the accumulation paths based on reaction times (RTs) and choices. (**c**) Schematic illustration of the hypothesized effects of increasing stimulus ambiguity on speed of evidence accumulation towards two types of responses. Each gray line represents the trajectory of evidence accumulation at a given level of perceptual ambiguity. As perceptual ambiguity increases, the evidence accumulates more slowly towards both boundaries (from *v*_1_ to *v*_n_), suggesting an interaction between ambiguity level and stimulus type. **(d)** Illustration of the interaction effect in the behavioral DDM models: increasing ambiguity is associated with slower evidence accumulation for both stimulus categories, yielding a negative ambiguity-by-stimulus category interaction on drift rate. **(e)** A hypothetical pattern of the posterior density distribution of the ambiguity × stimulus category (Type1 – Type0) interaction effect on drift rate. The x-axis shows the values of the parameter (e.g., the drift rate difference between fearful and happy facial stimuli), and the y-axis represents the probability density of each value based on the posterior distribution. **(f)** A previous EEG study using the same emotion categorization task showed that the magnitude of the Late Positive Potential (LPP) in the centroparietal area varied negatively with increasing perceptual ambiguity of the stimuli (37). **(g-i)** Neural model and posterior estimate of LPP × stimulus category interaction on drift rate. **(g)** Schematic illustration of the hypothesized effects of increasing LPP magnitude on speed of evidence accumulation towards two types of responses. As LPP amplitude increases, we hypothesized that the evidence accumulation towards fearful (Type 1) and happy (Type 0) responses would accelerate (from *v*_1_ to *v*_n_). **(h)** Illustration of the LPP × stimulus category interaction effect in the neural DDM models. **(i)** A hypothetical pattern of posterior density distribution of the LPP × stimulus category (Type1 – Type0) interaction effect on drift rate.

To comprehensively examine how evidence accumulation is modulated by key experimental manipulation (i.e., perceptual ambiguity) and neural signatures (e.g., LPP magnitude, neuronal firing rate), we employed a multimodal approach. We first applied the DDM to characterize evidence accumulation process at the behavioral level. To link these behavioral dynamics to neural substrates, we conducted a series of studies: four EEG experiments, each with an independent group of participants (Experiment 1: N = 16; Experiment 2a: N = 23; Experiment 2b: N = 11; Experiment 3: N = 15; Experiment 4: N = 32), one single-neuron recording study (Experiment 5, N = 16), and one fMRI study (Experiment 6, N = 19) (see **Table S1** for details). Together, this framework enables an integrated understanding of how evidence accumulation is neurally encoded during social-affective decision-making involving perceptual ambiguity.

### Ethical approval

Participants provided written informed consent according to protocols approved by the institutional review board and/or relevant authorities of the respective data collection site: South China Normal University (Experiments 1 – 4 (Sun, Zhen, et al., 2017), Experiment 6 (Sun, Yu, et al., 2023)) and Cedars-Sinai Medical Center (Experiment 5 (S. Wang et al., 2017)).

### Experiments 1 – 4: EEG Study

#### Participants

We recruited the following groups of participants for EEG experiments with different tasks (see **Table 1** for a summary). Experiment 1: 11 female and 5 male, mean age and SD: 19.6 ± 1.0 years. Experiment 2a: 17 female and 6 male, 22.4 ± 2.2 years. Experiment 2b/2c: 9 female/2 male, 20.6 ± 2.80 years. Experiment 3: 9 female and 6 male, 20.7 ± 1.61 years. Experiment 4: 15 female and 17 male, 20.6 ± 1.79 years. All participants had normal or corrected-to-normal visual acuity.

**Table 1.** Participant information across experiments. This table provides demographic and study-related information for all participants included in the analyses.

| Modality | Experiment Index | Stimulus | Task | Response type | Sample size (N) | Sex (M/F) | Age (mean ±SD years) | CR | Trials | Test location | Participants | Language |
| --- | --- | --- | --- | --- | --- | --- | --- | --- | --- | --- | --- | --- |
| EEG | Experiment 1a | Fear-Happy Morph | Fear-Happy (Emotion) Judgment | Speeded | 16 | 5/11 | 19.6 ± 0.96 | Y | 280 trials in 2 blocks | SCNU | University students | Chinese |
|  | Experiment 1b | Fear-Happy Morph | Male-Female (Gender) Judgment | Speeded |  |  |  |  |  |  |  |  |
|  | Experiment 1c | Fear-Happy Morph | Rich-Poor (Wealth) Judgment | Speeded |  |  |  |  |  |  |  |  |
| | Experiment 2a | Fear-Happy Morph | Fear-Happy Judgment | Non-speeded | 23 | 6/17 | 22.4 ± 2.17 | N | 252 trials in 2 blocks | SCNU | | |
| | Experiment 2b | Anger-Disgust Morph | Anger-Disgust Judgment | Non-speeded | 11 | 2/9 | 20.6 ± 2.80 | N | 252 trials in 2 blocks | SCNU | | |
|  | Experiment 2c | Cat-Dog Morph | Cat-Dog Judgment | Non-speeded |  |  |  | N | 252 trials in 2 blocks | SCNU |  |  |
| | Experiment 3 | Angry-Disgust/Cat-Dog Morph | Human-Animal judgment | Non-speeded | 15 | 6/9 | 20.7 ± 1.61 | N | 252 trials in 2 blocks | SCNU | | |
| | Experiment 4 | Fear-Happy Morph | Certain response | Non-speeded | 32 | 17/15 | 20.6 ± 1.79 | N | 192 trials in 3 blocks | SCNU | | |
| Single Neuron Recording | Experiment 5 | Fear-Happy Morph | Fear-Happy Judgment | Non-speeded | 16 | 11/5 | 42.3 ± 17.0 | Y | 176 to 440 trials in 2 to 5 blocks | CSMC | Neurosurgical patients | English |
| fMRI | Experiment 6 | Fear-Happy Morph | Fear-Happy Judgment | Non-speeded | 19 | 4/15 | 20.9 ± 2.02 | N | 168 trials in 2 blocks | SCNU | University students | Chinese |
| Total |  |  |  |  | 132 | 51/81 |  |  |  |  |  |  |

*Note:* CR indicates whether confidence ratings were included in the study. Abbreviation of institution names: SCNU = South China Normal University (Guangzhou, China), CSMC = Cedars-Sinai Medical Center (Los Angeles, U.S.). EEG data was adapted from Sun et al., 2017 NeuroImage; Sun et al., 2017, eNeuro. Single neuron recording data was adapted from Wang et al., 2017, Nature Communications. fMRI data was adapted from Sun et al., 2023, Translational Psychiatry

#### Experimental procedure

##### Experiment 1

Participants completed a perceptual decision-making task with fear-happy morphed face stimuli. Past research has shown that fear and happiness expressions are clearly distinguishable by facial features (Smith et al., 2005). We selected faces of four individuals (2 female, 2 male) each posing highly recognizable fear and happiness expressions from the STOIC database (Roy et al., 2010). Selected faces served as anchors and were the unambiguous exemplars of fearful and happy expressions. The database also provides normative ratings of valence and arousal of these facial stimuli. We then generated a series of morphed expression continua for this experiment by interpolating pixel values and locations between fearful and happy exemplar faces. Specifically, we applied a piecewise cubic-spline transformation over a Delaunay tessellation defined by manually selected control points placed on the main facial features (eyes, eyebrows, nose, and mouth). Control points were placed consistently across all anchors by two independent experimenters and were cross-checked for alignment accuracy, minimizing experimenter bias. We generated five levels of fear–happy morphs, ranging from 30% fear / 70% happy to 70% fear / 30% happy in steps of 10%. Together with the anchor images, there were seven stimuli for each face. Low-level image properties were equalized using the SHINE toolbox (Willenbockel et al., 2010). Independent subjective ratings confirmed that perceived ambiguity tracked the morph continuum (**Figure S1**), with the highest ambiguity near the categorical boundary, indicating that the stimuli captured genuine perceptual uncertainty rather than mere morphing artifacts. The complete set of continua and subjective ratings is shown in **Figure S1**.

This experiment consisted of three blocks, each with a different task instruction (**Figure 2a**). In the facial expression block (**Experiment 1a**), the participants judged whether a morphed face image displayed a fearful or happy expression. In the gender block (**Experiment 1b**), the participants judged whether the face image was of a male or a female individual. In the wealth block (**Experiment 1c**), the participants judged whether the face image was of a wealthy or a poor individual. The participants completed this task with speeded responses; namely, they could press a button to respond as soon as the stimulus was presented.

**Figure 2.**
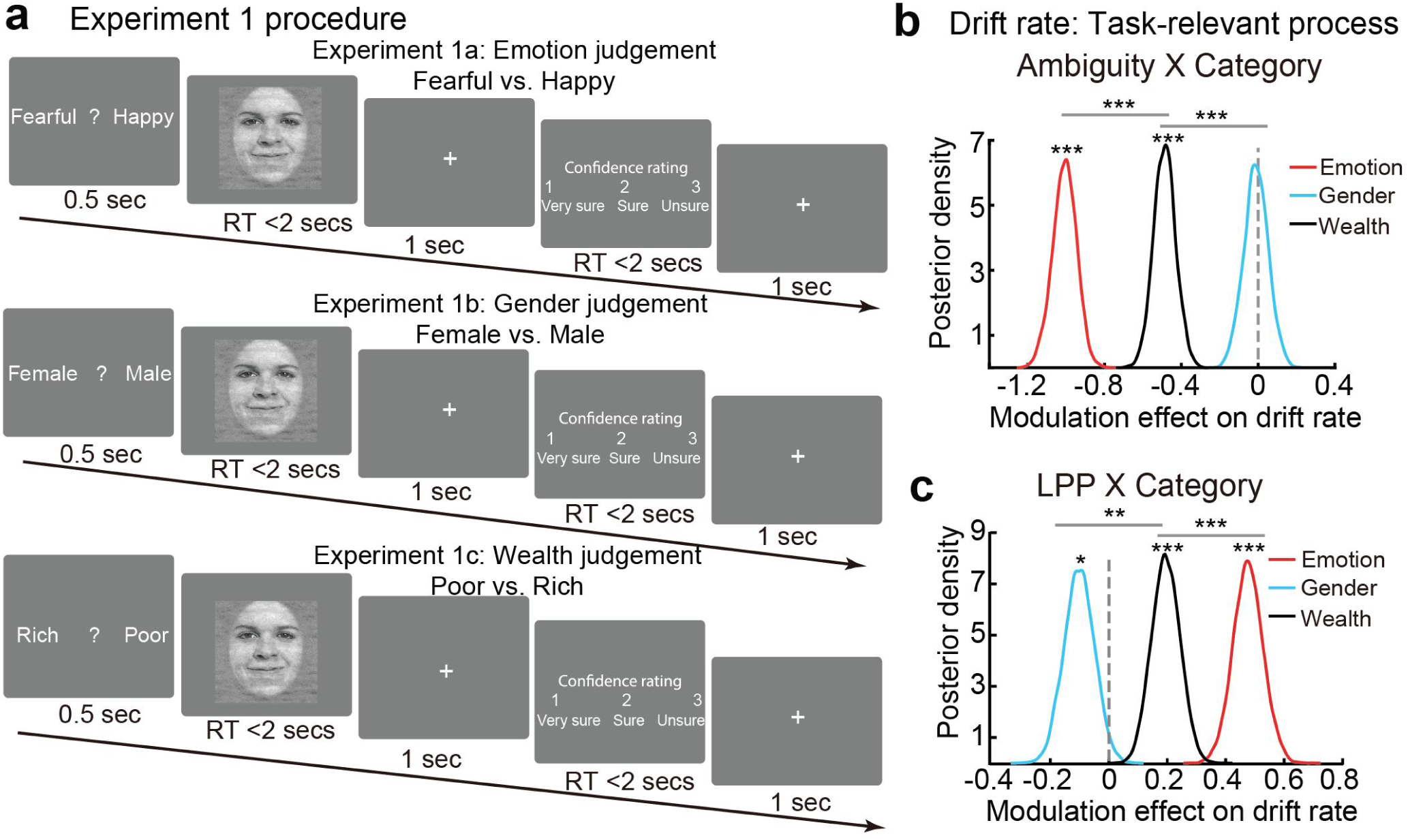
Task procedure and drift diffusion modeling results. **(a)** Experimental task: judgments based on different features of the same stimuli, i.e., emotion, gender, and wealth. Note that the facial images shown here are adopted from a publicly available database^63^. **(b)** A negative category × ambiguity modulation effect on drift rate was observed in the emotion judgment task, the effect size of which was significantly larger than that in the gender and wealth judgment tasks. **(c)** A positive category × LPP modulation effect on drift rate was observed in the emotion judgment task, the effect size of which is significantly larger than that in the gender and wealth judgment tasks. * Probability > 95%; ** Probability > 99%; *** Probability > 99.9%.

On each trial, participants were presented with a question prompt reminding them of the current task (expression, gender, wealth). The prompt was displayed for 500 ms, followed by a face stimulus presented for 2 seconds. Participants were instructed to respond as quickly as possible by pressing a button to indicate their judgment. The stimulus remained on the screen until a response was registered, and participants had up to 2 seconds to respond. If no response was registered within 2 seconds, the trial was aborted and excluded from data analysis. No feedback was provided during the task, and the order of facial stimuli was fully randomized across blocks and across participants. An inter-trial interval (ITI) with a central fixation cross was jittered randomly using a uniform distribution ranging from 1 to 2 seconds. To familiarize themselves with the task, participants completed 5 practice trials before the experiment began.

A subset of participants also provided confidence ratings. After the emotion judgment and a 500 ms blank screen, they were asked to indicate their confidence level by pressing one of three buttons: ‘1’ for ‘very sure,’ ‘2’ for ‘sure,’ or ‘3’ for ‘unsure.’ Participants had a maximum of two seconds to respond to the confidence question.

##### Experiment 2

As a replication and extension of **Experiment 1**, we recruited another two groups of participants to complete similar perceptual decision-making tasks. For **Experiment 2a**, participants completed the same fear-happy facial expression discrimination task as in the expression block of **Experiment 1** (**Figure 3a**). The only difference was that the participants completed this task with non-speeded responses, where the stimulus remained on the screen for 1 second, after which the participants pressed a button to respond. This way, participants viewed the stimuli for at least 1 second on every trial, giving us a cleaner measure of stimulus-induced late positive potential (LPP). All the remaining tasks reported in this paper adopted non-speeded responses. As with the speeded responses, the participants had up to two seconds to respond. The ITI and randomization protocols were identical to those in the main task.

**Figure 3.**
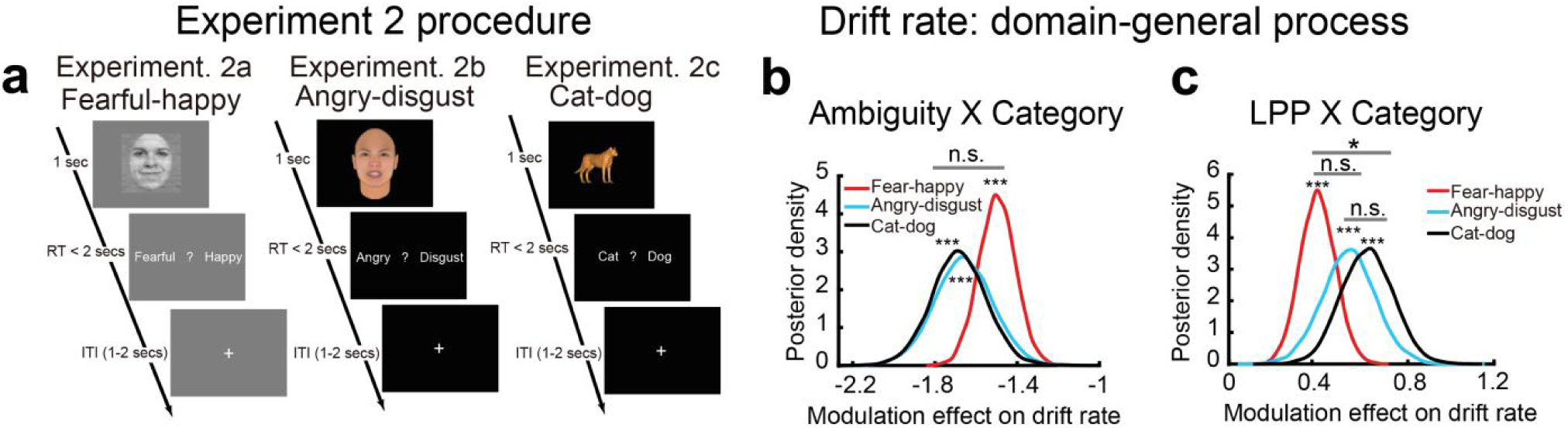
Task procedure and results of Experiment 2. **(a)** Domain-general task variants: perceptual judgments involving fearful-happy morphed facial images, angry-disgust morphed facial images, and cat-dog morphed images. Note that the facial images shown here are either adopted from a publicly available database^63^ (**Experiment 2a**) or computer-generated (**Experiment 2b**). **(b)** We identified negative category-by-ambiguity modulation effect on drift rates across the three tasks at the behavioral level. **(c)** We identified positive category × LPP modulation effect on drift rates at the neural level. * Probability > 95%; *** Probability > 99.9%.

**Experiment 2b** and **Experiment 2c** were conducted on the same group of participants. For **Experiment 2b**, the participants completed a perceptual decision-making task with different visual stimuli. Specifically, they viewed computer-generated anger-disgust morphed face images and judged whether an image expresses anger or disgust (**Figure 3a**). The face images were created using FaceGen Modeller (http://facegen.com/). We selected 4 individuals (2 male and 2 female) derived from 3D human face models, including two Asian (1 male, 1 female) and two Caucasian individuals (1 male, 1 female). Each individual had 2 anchor expressions (100% anger and 100% disgust) and 5 morph levels, ranging from 30% anger/70% disgust to 70% anger/30% disgust, with 10% increment. For the cat-dog morphed images (**Experiment 2c)**, two cat identities and two dog identities were selected. Each cat identity was morphed with a dog identity to create 4 morph lines. Each morph line included 2 anchor images and 5 morph levels (20% cat/80% dog, 40% cat/60% dog, 50% cat/50% dog, 60% cat/40% dog, and 80% cat/20% dog) (Freedman et al., 2001).

##### Experiment 3

To further replicate the modulation of task relevance on the neurocognitive processes underlying ambiguity resolution, we recruited another 15 participants to view the same computer-generated anger-disgust morphed images as in **Experiment 2b** and cat-dog morphed images as in **Experiment 2c**. Their task, however, was to judge whether an image depicted a human or an animal (**Figure 4a**). Thus, the perceptual ambiguity within each category (anger-disgust and dog-cat) was not relevant to the participants’ categorization tasks.

**Figure 4.**
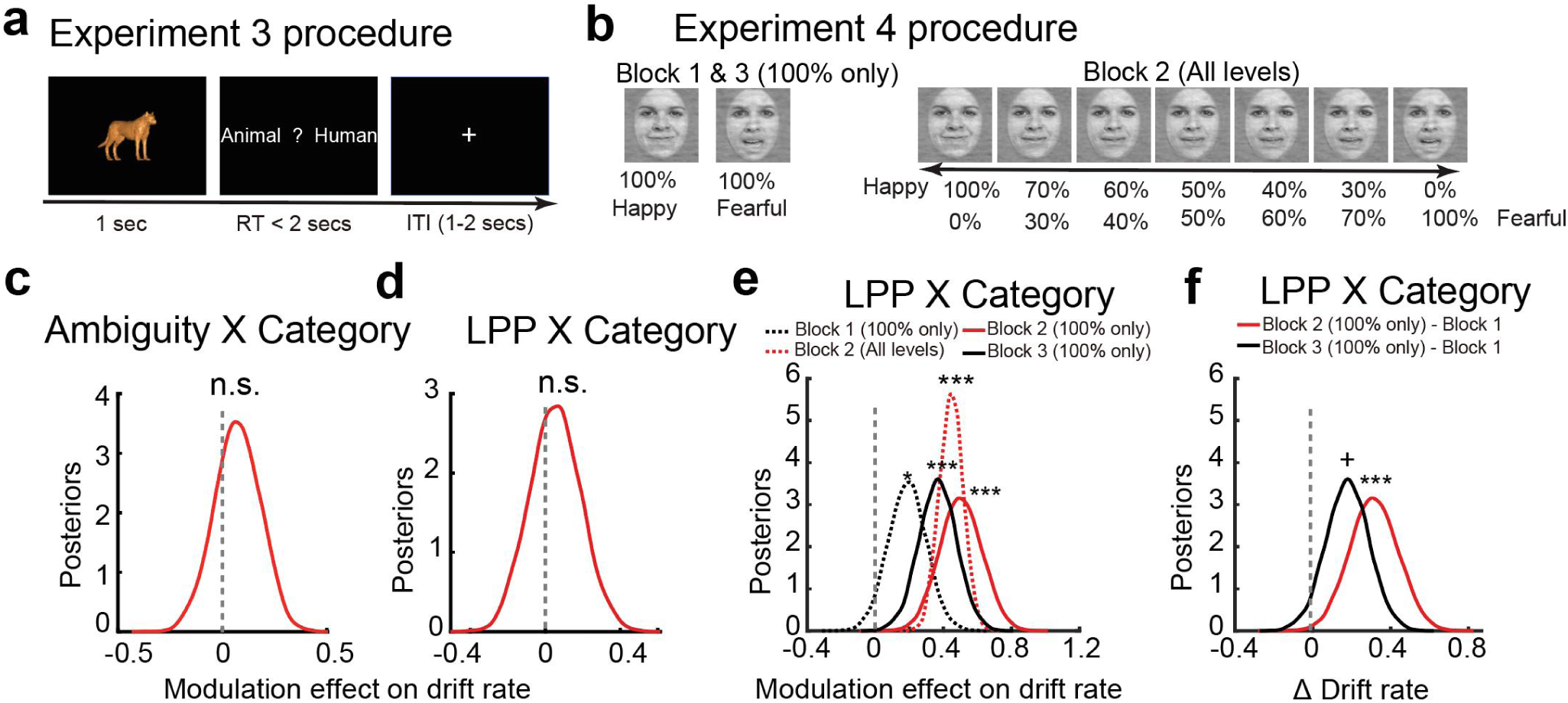
Task procedure and results of Experiment 3 and 4. **(a, b)** Task variants with contextual modulation. Note that the facial images shown here (**b**) are adopted from a publicly available database^63^. **(c, d)** Decisions made under perceptual ambiguity following certain contexts did not show evidence accumulation, as reflected by non-significant modulation effect on drift rates at both the **(c)** behavioral and **(d)** neural levels. (**e**) Category × LPP modulation on drift rates were significantly reduced in Block 1 (100% certainty levels) compared to Block 2 (mixed ambiguity) and Block 3 (later repetition). **(f)** Differences in drift rates between blocks, with Blocks 2 and 3 showing significantly stronger drift rates compared to Block 1. + Probability > 90%; * Probability > 95%; *** Probability > 99.9%.

##### Experiment 4

This experiment consisted of three blocks (**Figure 4b**). In the middle block (Block 2), participants viewed the same fearful-happy morphed face images and were instructed to make fearful-happy binary choices as in **Experiment 1**. However, in Block 1 and Block 3, participants viewed and judged only unambiguous faces (i.e., the anchor faces).

#### EEG data acquisition and analysis

Procedures for EEG data recording and analyses were described in detail in our previous work (Sun, Yu, et al., 2017a; Sun, Zhen, et al., 2017). Briefly, EEGs were recorded using a digital AC amplifier from 32 scalp sites with electrodes mounted in an elastic cap (NeuroScan 4.5) according to the International 10–20 system. EEGs were recorded from the following sites: frontal: FP1, FP2, F7, F3, Fz, F4, F8; frontal-central: FC3, FCz, FC4; central: C3, Cz, C4; central-parietal: CP3, CPz, CP4; parietal: P7, P3, Pz, P4, P8; frontal-temporal-parietal: FT7, TP7, T7, T8, TP8, FT8; and occipital: O1, Oz, O2. The ground electrode was placed on the forehead, and all recordings were referenced to the right mastoid. All impedance was maintained below 5 KΩ. EEG and electro-oculogram (EOG) were amplified using a 0.05-70 Hz band-pass filter and were continuously sampled at 500 Hz/channel. EEG data were processed using EEGLAB (84). The continuous EEG data were re-referenced to the average of the left and right external mastoid signals. The data were filtered using a digital zero-phase shift band-pass filter of 0.5-30 Hz with a slope of 24 dB/octave. Then the continuous EEG data were epoched into 1-second segments (–200 to 800 ms relative to stimulus onset), and the pre-stimulus interval (–200 to 0 ms) was used as the baseline. The data were then baseline-corrected by subtracting the average activity during the baseline period. Trials that had blinks or saccades were excluded, and the remaining artifacts were further detected using a moving-window peak-to-peak artifact detection method on specific midline electrodes. Within each participant, the mean waveform of each morph level was computed, time-locked to the onset of the stimulus. Single-participant mean waveforms were subsequently averaged to obtain group-level mean waveforms. Here, we measured the LPP (entire waveform) based on the time window of 400 to 700 ms after stimulus onset at the parietal-central (Pz) electrode (Sabatinelli et al., 2006). Importantly, the scalp topography of the difference waveform between high ambiguity and anchor showed the most pronounced difference at Pz in this time window (Sun, Zhen, et al., 2017). These procedures were the same for all the studies reported in this paper.

#### Drift diffusion model (DDM)

We fit and compared several models predicting participants’ choices and reaction times using the Hierarchical Drift Diffusion Models (HDDM) (Pan et al., 2025; Wiecki et al., 2013). In all these models, we included four model parameters: drift rate (*v*), threshold (*a*), non-decision time (*t*), and initial bias (*z*). These parameters were estimated hierarchically at individual and group levels (**Figure 1b**). We used the HDDM regression function to estimate the potential effects of experimental manipulation and stimulus properties (e.g., category and perceptual ambiguity) on DDM parameters.

##### Behavioral models

We first fit and compared several behavioral models. In Model 1, the drift rate parameter was modulated only by the objective ambiguity level of the stimuli (1 = 100% fearful and 100% happy stimuli, 2 = 70% fearful and 70% happy stimuli, 3 = 60% fearful and 60% happy stimuli; the 50% fearful/happy stimuli were excluded because they do not objectively belong to either category). In Model 2, the drift rate parameter was modulated by both the objective stimulus category and the objective ambiguity level in an additive manner. Model 3 only included the interaction term between the stimulus category and the objective ambiguity. Model 4 included the two main effect terms and the interaction term. We did not hypothesize that trial-by-trial experimental manipulation (i.e., ambiguity levels) would modulate the other DDM parameters.

*Drift rate ∼ 1+ Ambiguity (Model 1)*
*Drift rate ∼ 1+ Category + Ambiguity (Model 2)*
*Drift rate ∼ 1+ Category : Ambiguity (Model 3)*
*Drift rate ∼ 1+ Category + Ambiguity + Category : Ambiguity (Model 4)*

We first evaluated the model using behavioral choices and response times from a speeded decision task (i.e., **Experiment 1a**). Ambiguity was coded as three objective levels, as described above. Following HDDM convention, RTs shorter than 50 ms or longer than 2 s were excluded. Less than 5% of the trials were excluded this way, comparable to previous literature using HDDM(Ratcliff & Tuerlinckx, 2002; Wiecki et al., 2013). All models were estimated using Markov Chain Monte Carlo (MCMC) sampling with 10,000 iterations, including 1,000 burn-in samples (i.e., the first 1,000 iterations were discarded), resulting in 9,000 posterior samples. We examined posterior distributions to evaluate the significance of decision parameters and their modulation by experimental manipulation.

We compared competing models using the Deviance Information Criterion (DIC), a standard metric for assessing Bayesian models, particularly hierarchical ones such as HDDM, based on the trade-off between model fit and complexity. Model 4 yielded the lowest DIC, indicating the best overall fit (see **Table 2**). To assess convergence in MCMC sampling, we examined posterior traces and autocorrelations, and reported the Gelman-Rubin statistic (R̂) in **Table 3**. All R̂ values were below 1.01, indicating excellent convergence of the MCMC sampling. The posterior predictive RTs and their comparison with observed RTs for each subject, based on the best-fitting model, are shown in **Figure S4**. Overall, the model demonstrates good recovery of RT distributions, with simulated RTs closely aligned with the observed data.

**Table 2.** Model comparison and selection based on Deviance Information Criterion (DIC) values.

| Modality | Models | DIC value |
| --- | --- | --- |
| Behavioral model<br>(Experiment 1a) | *Drift rate ~ 1+ Ambiguity (Model 1)* | 6250 |
|  | *Drift rate ~ 1+ Category + Ambiguity (Model 2)* | 4952 |
|  | *Drift rate ~ 1+ Category : Ambiguity (Model 3)* | 5887 |
|  | ***Drift rate ~ 1+ Category + Ambiguity + Category : Ambiguity (Model 4)*** | 4693 |
| EEG model<br>(Experiment 1a) | *Drift rate ~ 1+ LPP (Model 5)* | 6245 |
|  | *Drift rate ~ 1+ Category + LPP (Model 6)* | 4951 |
|  | *Drift rate ~ 1+ Category : LPP (Model 7)* | 6229 |
|  | ***Drift rate ~ 1+ Category + LPP + Category : LPP (Model 8)*** | 4867 |
| Amygdala neurons<br>(Experiment 5) | *Drift rate ~ 1+ Firing rate (Model 9)* | 3698 |
|  | *Drift rate ~ 1+ Category + Firing rate (Model 10)* | 4517 |
|  | *Drift rate ~ 1+ Category : Firing rate (Model 11)* | 3897 |
|  | ***Drift rate ~ 1+ Category + Firing rate + Category : Firing rate (Model 12)*** | 3687 |
| dmPFC neurons<br>(Experiment 5) | *Drift rate ~ 1+ Firing rate (Model 9)* | 1900 |
|  | *Drift rate ~ 1+ Category + Firing rate (Model 10)* | 2345 |
|  | *Drift rate ~ 1+ Category : Firing rate (Model 11)* | 2152 |
|  | ***Drift rate ~ 1+ Category + Firing rate + Category : Firing rate (Model 12)*** | 1887 |

*Note:* This table summarizes the DIC values for competing models across different modalities. The full model consistently exhibited the lowest DIC, indicating the best trade-off between model fit and complexity. Based on this criterion, the full model was selected as the best-fitting model and was used in all subsequent analyses.

**Table 3.** R̂ (Rhat) and Effective Sample Size (ESS) values for the best-fitting model.

| Modality | Main models | Effects | $\widehat{R}$ value | ESS value |
| --- | --- | --- | --- | --- |
| Behavioral model<br>(Experiment 1a) | $Drift\ rate \sim 1 + Category + Ambiguity + Category : Ambiguity$ (Model 4) | v_Category | 1.003 | 665.2 |
|  |  | v_Ambiguity | 1.004 | 674.0 |
|  |  | v_Category X Ambiguity | 1.003 | 741.9 |
| EEG model<br>(Experiment 1a) | $Drift\ rate \sim 1 + Category + LPP + Category : LPP$ (Model 8) | v_Category | 1.002 | 1493 |
|  |  | v_Ambiguity | 1.002 | 1639 |
|  |  | v_Category X Ambiguity | 1.002 | 1552 |
| Amygdala neurons<br>(Experiment 5) | $Drift\ rate \sim 1 + Category + Firing\ rate + Category : Firing\ rate$ (Model 12) | v_Category | 1.002 | 2638 |
|  |  | v_Ambiguity | 1.001 | 2854 |
|  |  | v_Category X Ambiguity | 1.003 | 1759 |
| dmPFC neurons<br>(Experiment 5) | $Drift\ rate \sim 1 + Category + Firing\ rate + Category : Firing\ rate$ (Model 12) | v_Category | 1.002 | 2586 |
|  |  | v_Ambiguity | 1.001 | 2407 |
|  |  | v_Category X Ambiguity | 1.001 | 3869 |

Note: All $\hat{R}$ values are below 1.01, indicating good convergence. All ESS values exceed 600, suggesting that the posterior estimates are reliable and based on effectively independent samples.

We then replicated the results using an independent dataset with non-speeded responses (**Experiment 2a**). In this task, an additional 1 second (corresponding to the stimulus presentation duration) was added to the original RTs. The same modeling procedure was applied as in **Experiment 1a**. Similarly, Model 4 yielded the lowest DIC in the non-speeded response data. Regression results based on Model 4 were comparable between **Experiment 1** and **Experiment 2a**, suggesting that the response mode did not substantially affect the reliability of the modeling results. In a supplementary analysis, ambiguity was also coded based on subjective ratings from an independent group of participants (**Figures S3**). Overall, we observed a qualitatively similar negative interaction of category-by-ambiguity modulation on drift rate. Based on these findings, we adopted the same model structure (Model 4), using objective ambiguity levels, for all subsequent DDM analyses. Summary statistics, including main effects and interactions for the best-fitting models, are presented in **Table S1**.

##### Neural models

Next, we incorporated the normalized, trial-by-trial LPP amplitude in the DDM models. Since we hypothesized that LPP amplitude is a neural signature of ambiguity resolution, we replaced the ambiguity level in the behavioral models with LPP amplitude. If the LPP indeed reflects a neural process that resolves perceived ambiguity, then the LPP should have the opposite effect of the ambiguity level. We estimated and compared four models similar to the four behavioral models, with the exception that the ambiguity level was replaced by LPP magnitude, as shown below.

*Drift rate ∼ 1+ LPP (Model 5)*
*Drift rate ∼ 1+ Category + LPP (Model 6)*
*Drift rate ∼ 1+ Category : LPP (Model 7)*
*Drift rate ∼ 1+ Category + LPP + Category : LPP (**Model 8**)*

All models were estimated using MCMC sampling with 10,000 iterations, including 1,000 burn-in samples (i.e., the first 1,000 iterations were discarded), resulting in 9,000 posterior samples. The full model (Model 8) had the lowest DIC, indicating that it was the best-fitting model (see **Table 2**). To assess convergence in MCMC sampling, we examined posterior traces and autocorrelations, and reported the Gelman-Rubin statistic (R̂) in **Table 3**. All R̂ values were below 1.01, indicating excellent convergence of the MCMC sampling. Our statistical inferences were then based on the posterior distributions of the best-fitting models. The model parameters of the best-fitting models are shown in **Figure 2** through **Figure 4**. Summary statistics, including main effects and interactions for the best-fitting models, are presented in **Table 4**.

**Table 4.**
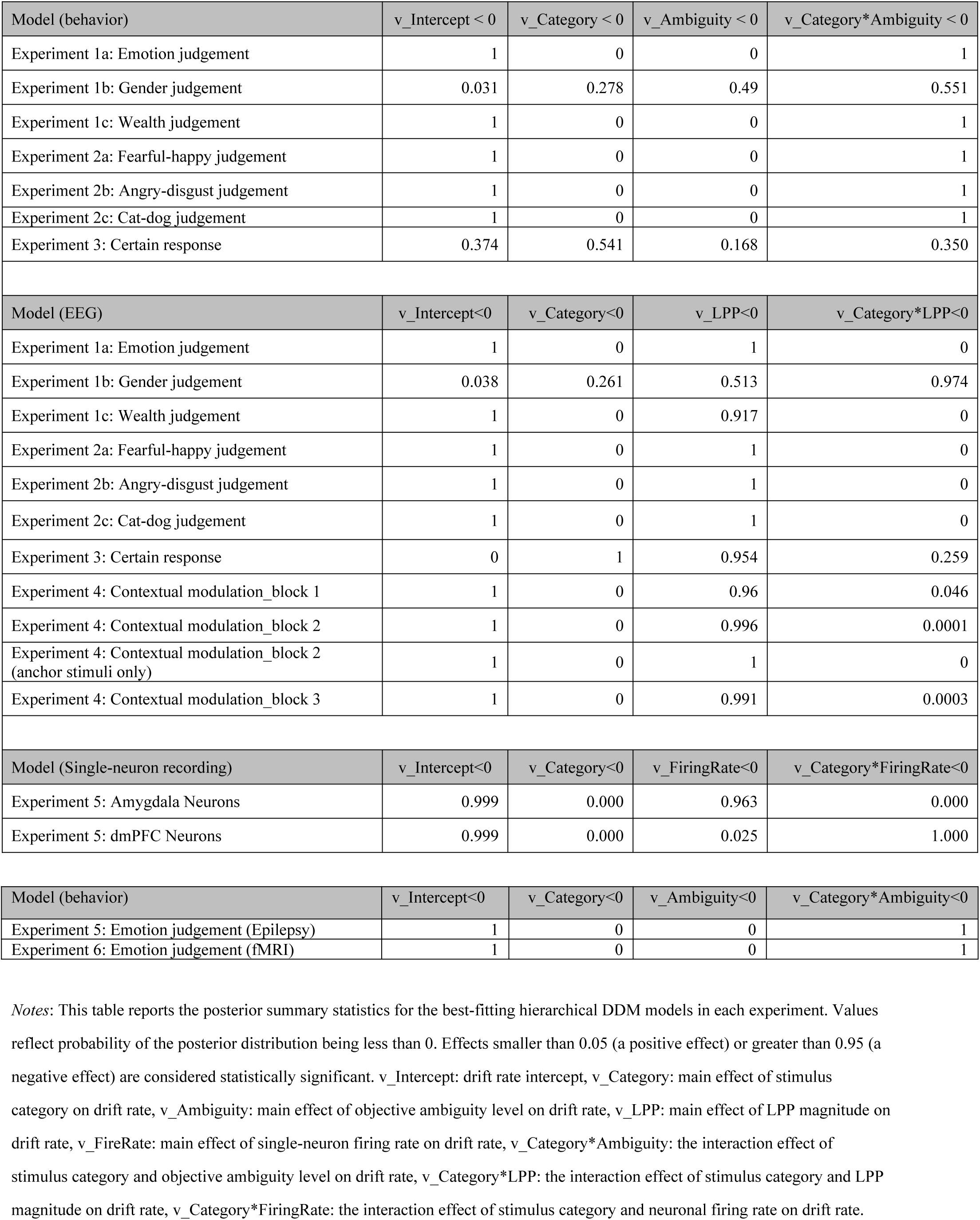
Parameter estimates from the winning models across all experiments.

| Model (behavior) | v_Intercept < 0 | v_Category < 0 | v_Ambiguity < 0 | v_Category*Ambiguity < 0 |
| --- | --- | --- | --- | --- |
| Experiment 1a: Emotion judgement | 1 | 0 | 0 | 1 |
| Experiment 1b: Gender judgement | 0.031 | 0.278 | 0.49 | 0.551 |
| Experiment 1c: Wealth judgement | 1 | 0 | 0 | 1 |
| Experiment 2a: Fearful-happy judgement | 1 | 0 | 0 | 1 |
| Experiment 2b: Angry-disgust judgement | 1 | 0 | 0 | 1 |
| Experiment 2c: Cat-dog judgement | 1 | 0 | 0 | 1 |
| Experiment 3: Certain response | 0.374 | 0.541 | 0.168 | 0.350 |
| Model (EEG) | v_Intercept<0 | v_Category<0 | v_LPP<0 | v_Category*LPP<0 |
| Experiment 1a: Emotion judgement | 1 | 0 | 1 | 0 |
| Experiment 1b: Gender judgement | 0.038 | 0.261 | 0.513 | 0.974 |
| Experiment 1c: Wealth judgement | 1 | 0 | 0.917 | 0 |
| Experiment 2a: Fearful-happy judgement | 1 | 0 | 1 | 0 |
| Experiment 2b: Angry-disgust judgement | 1 | 0 | 1 | 0 |
| Experiment 2c: Cat-dog judgement | 1 | 0 | 1 | 0 |
| Experiment 3: Certain response | 0 | 1 | 0.954 | 0.259 |
| Experiment 4: Contextual modulation_block 1 | 1 | 0 | 0.96 | 0.046 |
| Experiment 4: Contextual modulation_block 2 | 1 | 0 | 0.996 | 0.0001 |
| Experiment 4: Contextual modulation_block 2 (anchor stimuli only) | 1 | 0 | 1 | 0 |
| Experiment 4: Contextual modulation_block 3 | 1 | 0 | 0.991 | 0.0003 |
| Model (Single-neuron recording) | v_Intercept<0 | v_Category<0 | v_FiringRate<0 | v_Category*FiringRate<0 |
| Experiment 5: Amygdala Neurons | 0.999 | 0.000 | 0.963 | 0.000 |
| Experiment 5: dmPFC Neurons | 0.999 | 0.000 | 0.025 | 1.000 |
| Model (behavior) | v_Intercept<0 | v_Category<0 | v_Ambiguity<0 | v_Category*Ambiguity<0 |
| Experiment 5: Emotion judgement (Epilepsy) | 1 | 0 | 0 | 1 |
| Experiment 6: Emotion judgement (fMRI) | 1 | 0 | 0 | 1 |

*Notes:* This table reports the posterior summary statistics for the best-fitting hierarchical DDM models in each experiment. Values reflect probability of the posterior distribution being less than 0. Effects smaller than 0.05 (a positive effect) or greater than 0.95 (a negative effect) are considered statistically significant. v\_Intercept: drift rate intercept, v\_Category: main effect of stimulus category on drift rate, v\_Ambiguity: main effect of objective ambiguity level on drift rate, v\_LPP: main effect of LPP magnitude on drift rate, v\_FireRate: main effect of single-neuron firing rate on drift rate, v\_Category\*Ambiguity: the interaction effect of stimulus category and objective ambiguity level on drift rate, v\_Category\*LPP: the interaction effect of stimulus category and LPP magnitude on drift rate, v\_Category\*FiringRate: the interaction effect of stimulus category and neuronal firing rate on drift rate.

### Experiment 5: Single-unit Recording Study

#### Participants

Nine neurosurgical patients (5 female/4 male, 42.3 ± 17.0 years) completed fourteen recording sessions (three patients completed two sessions, and one patient completed three sessions). Each session was treated as an independent sample for behavioral analysis. All participants had normal or corrected-to-normal visual acuity.

#### Single-neuron electrophysiology recording

We recorded neurons from implanted depth electrodes in the amygdala and dmPFC in patients with pharmacologically intractable epilepsy. Target locations were verified using post-implantation CT. At each site, we recorded from eight 40 μm microwires inserted into a clinical electrode as described previously(Rutishauser, Mamelak, et al., 2006; S. Wang et al., 2014). Bipolar wide-band recordings (0.1-9000 Hz), using one of the eight microwires as the reference, were sampled at 32 kHz and stored continuously for off-line analysis with a Neuralynx system. The raw signal was filtered with a zero-phase lag 300-3000 Hz bandpass filter, and spikes were sorted using a semi-automatic template matching algorithm, as described previously (Rutishauser, Schuman, et al., 2006). Units were carefully isolated and recording and spike sorting quality were assessed quantitatively. Only units with an average firing rate of at least 0.2 Hz (entire task) were considered. We quantified the response of each neuron based on the number of spikes observed after stimulus onset (1.5 s window, starting 250 ms after stimulus onset). Full details of the single-neuron recording methods (including the number of recorded units per region, inclusion criteria, and variability across participants), has been reported in our previous publications (S. Wang et al., 2017). In the current manuscript, we provide a summary of the key parameters for clarity, and the relevant summary statistics are included in **Table S1**.

#### Spikes

Only units with an average firing rate of at least 0.2 Hz (entire task) were considered. Only single units were considered. Trials were aligned to face onset, and the baseline firing rate was calculated in a 1 second interval of a blank screen immediately before face onset. Average firing rates and Peri-Stimulus Time Histogram (PSTH) were computed by counting spikes across all trials in consecutive 250 ms bins. Comparisons between morph levels were made using a one-way ANOVA at P < 0.05 and Bonferroni-corrected for multiple comparisons in the group PSTH.

#### DDM

We first replicated the behavioral DDM model identified in Study 1, where the drift rate was modulated by stimulus category, ambiguity level, and their interaction. We next applied the same neural DDM model as in Study 1 to investigate how the DDM parameters were encoded by neuronal activities. Specifically, we extracted the firing rate from the bilateral amygdala and dmPFC individually and examined how the drift rate was modulated by stimulus categories, neuronal firing rates, and their interaction on a trial-by-trial basis:

*Drift rate ∼ 1+ Firing Rate (Model 9)*
*Drift rate ∼ 1+ Category + Firing Rate (Model 10)*
*Drift rate ∼ 1+ Category : Firing Rate (Model 11)*
*Drift rate ∼ 1+ Category + Firing Rate + Category : Firing Rate (**Model 12**)*

As in Experiment 2a, RTs faster than 50 ms and longer than 2 seconds were discarded from the analysis. Trials excluded due to this criterion were less than 5% of the total trials. An extra 1 sec (the presentation duration of the stimuli) was added to the original RTs because participants could only respond after the stimulus presentation. In addition, trials with the highest ambiguity (50% morphed levels) were excluded because they could not be objectively classified into either category. All models were estimated using MCMC sampling with 10,000 iterations, including 1,000 burn-in samples (i.e., the first 1,000 iterations were discarded), resulting in 9,000 posterior samples. The full model (***Model 12***) had the lowest DIC score, indicating that it was the best-fitting model (see **Table 2**). To assess convergence in MCMC sampling, we examined posterior traces and autocorrelations, and reported the Gelman-Rubin statistic (R̂) in **Table 3**. All R̂ values were below 1.01, indicating excellent convergence of the MCMC sampling.

Parameter estimates were derived from the posterior distributions of the best-fitting model (***Model 12***). The parameters of the best-fitting model are shown in **Figure 5f** and **5g**. Summary statistics, including main effects and interactions for the best-fitting models, are presented in **Table S4**. For comparison, we also tested reduced models, including one with only the main effect of firing rate (***Model 9***), one with the main effects of both stimulus category and firing rate (***Model 10***), and one with the interaction between category and firing rate (***Model 11***). The performance of these models was worse than the best-fitting model (see **Table 2** for model comparison details).

**Figure 5.**
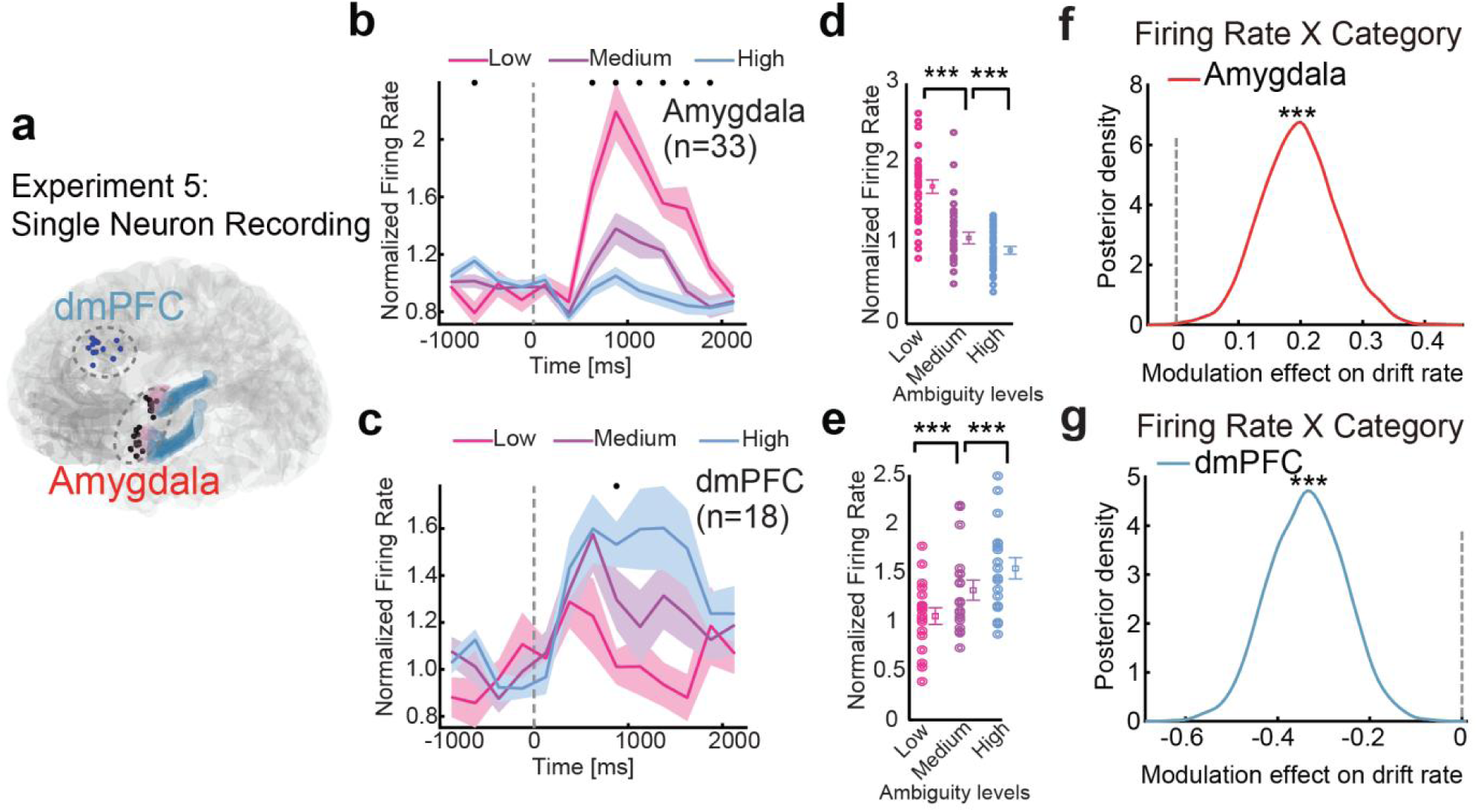
Single-neuron correlates of decision parameters (Experiment 5). **(a)** Neurons recorded from the dmPFC and amygdala sites. (**b-c**) Average normalized firing rate of ambiguity-coding neurons. Asterisk indicates a significant difference between the conditions in that bin (P < 0.05, one-way ANOVA, Bonferroni-corrected). **(d-e)** Mean normalized firing rate at ambiguity level. Normalized firing rate for each unit (left) and mean ± SEM across units (right) are shown at each ambiguity level. Mean firing rate was calculated in a time window 250–1750 ms after stimulus onset (the same time window as neuron selections). Asterisks indicate a significant difference between conditions using paired two-tailed t-test. *** P < 0.001. **(b, d)** Neurons in the amygdala that increased their firing rate for the least ambiguous faces (*n* = 33). **(c, e)** Neurons in the dmPFC that increased their firing rate for the most ambiguous faces (*n* = 18). **(f)** Amygdala neurons exhibited a positive category × firing rate interaction effect on drift rate. **(g)** dmPFC neurons exhibited a negative category × firing rate interaction effect on drift rate. *** Probability > 99.9% (in panels **f** and **g**).

### Experiment 6: fMRI Study

#### Participants

Nineteen healthy, right-handed volunteers (15 female/4 male, 20.9 ± 2.02 years) participated in the fMRI experiments with fear-happy non-speeded judgments. All participants had normal or corrected-to-normal visual acuity.

#### Experimental design and procedure

The experimental design is similar to the main non-speeded fear-happy judgment task in **Experiment 2a** (**Figure 3a**), except that the ITI was jittered randomly with a uniform distribution between 2–8 seconds. The procedure was described in detail elsewhere (Sun, Zhen, et al., 2017; S. Wang et al., 2017).

#### DDM

We applied the same best-fitting behavioral model (***Model 4***) in the EEG study to the behavioral data from the fMRI study to investigate how decision variables are encoded at the neural circuit level. To model the drift rate, we used the same HDDM regression model as in **Experiment 2a**, incorporating the main effects of stimulus category and ambiguity level, as well as their interaction. All modeling criteria were identical to those used for the EEG participants, with one key difference: we additionally estimated each participant’s drift rate as a function of the category-by-ambiguity interaction. From this model, we extracted individual drift rate estimates for the category-by-ambiguity interaction to examine their association with frontal – amygdala connectivity strength. Posterior estimates of the model are reported in **Table S4**, which show a similar pattern as the behavioral model of **Experiment 2a**.

#### fMRI acquisition and preprocessing

Procedures for fMRI data acquisition and analyses were described in detail in our previous studies (37, 83). Briefly, MRI scanning was conducted at South China Normal University on a 3-Tesla Tim Trio Magnetic Resonance Imaging scanner (Siemens, Germany) using a standard 12-channel head-coil system. Whole-brain data were acquired with echo planar T2*-weighted imaging (EPI), sensitive to blood-oxygen-level dependent (BOLD) signal contrast (31 oblique axial slices, 3 mm thickness; TR = 2000 ms; TE = 30 ms; flip angle = 90°; FOV = 224 mm; voxel size: 3 × 3 × 3 mm). T1 weighted structural images were acquired at a resolution of 1 × 1 × 1 mm. Neuroimaging data were preprocessed using SPM8, including slice-timing correction, realignment, smoothing with a Gaussian kernel of full-width-half-maximum 6 mm, and co-registration of EPI and structural images to the T1 MNI 152 template. To provide inputs to the DDM, we fitted a GLM at face onset and derived a beta value for each face. We averaged the beta values across participants to get a mean beta value for each face and each voxel. Full details of the fMRI acquisition parameters, preprocessing steps, and statistical modeling procedures have also been reported in our previous publications (Sun, Zhen, et al., 2017; Sun, Cao, et al., 2023; Sun, Yu, et al., 2023; S. Wang et al., 2017).

#### fMRI functional connectivity analysis

Our previous work has shown that the dmPFC exerts top-down modulation on the right amygdala during decisions under ambiguity. In addition, the connectivity strength between the dmPFC and the right amygdala, characterized by Psychophysiological Interaction model (PPI), was positively modulated by individual participants’ sensitivity to ambiguity, as measured by reaction time differences between high and low ambiguity levels (Sun, Yu, et al., 2023), suggesting that the dmPFC-amygdala could be a target circuit in representing decision parameters dependency on ambiguity levels. Given that our main goal is to identify whether and how the decision parameters, particularly the drift rates, are encoded by the neural circuit, we extracted the dmPFC-right amygdala connection strength from the PPI model and correlated it with individual drift rates derived from the DDM. We hypothesized that the dmPFC-right amygdala connectivity strength would be negatively correlated with the category-by-ambiguity modulation on drift rate derived from the DDM.

## Results

### Drift diffusion models capture both behavioral and neural correlates of evidence accumulation

Previous research has shown that drift rate (*v*) is associated with evidence accumulation processes in sensorimotor and economic decision-making contexts and plays a key role in differentiating decisions based on varying levels of ambiguity or difficulty (Hadar et al., 2016; M. A. Pisauro et al., 2017; Polanía et al., 2014). Based on this, we hypothesized that drift rate (*v*) may serve as a key decision parameter that reflects how ambiguity level modulates evidence accumulation on a trial-by-trial basis; this mechanism should be generalizable across different domains of categorization decisions (e.g., emotion categories vs. animal species) under perceptual ambiguity.

To test this hypothesis, we applied identical behavioral and neural DDM models to the main task (**Figure 1a**), which involved categorization decisions for fear-happy morphed faces (**Experiment 1, Experiment 2a**), as well as to several task variants using different stimulus categories (**Experiment 2b**: angry-disgust morphed faces; **Experiment 2c**: cat-dog morphed images) (**Figure 2a**; **Figure 3a**) and the sensitivity of these effects across decision contexts (**Experiment 3 and 4**: **Figures 4a, 4b**). We included the following four DDM parameters in all models: drift rate, initial bias, threshold, and non-decision time (**Figure 1b**). Based on our prior observations that reaction times (RTs) increase with increasing ambiguity level, we hypothesized that the speed of evidence accumulation (i.e., drift rate | U |) would decrease with increasing ambiguity level for both category boundaries, leading to a negative ambiguity × stimulus category interaction effect on drift rate (**Figures 1c – 1e**). Since stimulus category and ambiguity levels were transient, varying in a trial-by-trial manner, we did not hypothesize that they would influence decision threshold or initial bias.

Neurally, previous work has identified the LPP as a neural marker that discriminates ambiguity levels (see **Figure 1f**) (Sun, Yu, et al., 2017a; Sun, Zhen, et al., 2017). For example, it has been shown that it builds gradually with sensory evidence and reaches a threshold at response execution, independent of sensory modality or motor processes (O’Connell et al., 2012; O’Connell & Kelly, 2021). We therefore hypothesized that as LPP amplitude increases, participants’ percepts should be less ambiguous and the evidence accumulation towards either boundary should be faster, leading to a positive LPP × stimulus category interaction effect on drift rate (**Figures 1g – 1i**). We estimated and compared several models that are different in terms of whether stimulus category, ambiguity level (for behavioral models), LPP magnitude (for neural models), and/or their interaction modulate drift rate (**Table 2**). In the winning model, drift rate is modulated by stimulus category, perceptual ambiguity level, and their interaction. We then examined the posterior distribution of the parameters to test our hypotheses.

**Experiment 1** consisted of three blocks, during which the participants viewed the same fearful-happy morph images. However, each block had a different task instruction. In the facial expression block, the participants judged whether a morphed face image was a fearful or a happy expression. In the gender block, the participants judged whether the face image was a male or a female person. In the wealth block, the participants judged whether the face image was a wealthy or a poor person. The participants completed this task with speeded responses; namely, they could press a button to respond as soon as the stimulus was presented. As predicted, we observed a significant negative category × ambiguity interaction effect on drift rate in the facial expression judgment block (99.9%), but no such effect was observed in the gender judgment block (55.1%) (**Figure 2b**). Interestingly, we also observed a significant negative category × ambiguity interaction effect on drift rate in the wealth judgment task (99.9%). While this was not initially predicted, it was in line with the literature showing that positive facial emotion biases the judgment of wealth and social status (Galinsky et al., 2020; Todorov, 2008). It was also consistent with the behavioral pattern in our dataset, where we found that happier faces were reliably judged as wealthier and faces with clear emotional expressions elicited higher confidence in wealth judgments (**Figures S9e, S9j, S9l**). More importantly, the effect size of the interaction was significantly larger in the facial expression judgment block than in the other two blocks (all posterior probabilities > 95%). This indicates that perceptual ambiguity hinders evidence accumulation primarily when the ambiguity is task-relevant. This pattern also mirrors the behavioral ambiguity modulation shown in **Figures S1** and **S2**. Subjective measures (perceived ambiguity, and confidence ratings) and objective measures (RTs) both showed a graded ambiguity interference effect across the three tasks (facial expression task > wealth > gender).

To enhance transparency in our modeling results, we additionally visualized the posterior probability densities of the individually fitted drift rates across stimulus categories and ambiguity levels (**Figure S3b**, **Table S2**). These plots allow direct inspection of how evidence accumulation strength varies with perceptual ambiguity. For the facial emotion judgment task (both speeded and non-speeded versions), the posterior distributions for the least ambiguous stimuli (anchor faces) are shifted furthest toward positive and negative drift rates for the two categories, indicating strong, unambiguous accumulation toward the respective decision boundaries **(Figures S3b, S3c)**. In contrast, the distributions for the most ambiguous stimuli cluster near zero, reflecting weaker accumulation. This pattern provides an intuitive illustration of the stimulus category × ambiguity interaction in the winning models reported above and confirms that drift rate systematically captures the impact of perceptual ambiguity on decision formation. Lastly, this negative category × ambiguity interaction effect was further replicated in a non-speeded version of the task (**Experiment 2)**, where the participants could only press a button to register their response one second after the stimulus onset (**Figure S3a, S3c**).

As a robustness check, in a control analysis, we replaced researcher-defined objective ambiguity (i.e., morph level) with subjective ambiguity rated by an independent group of participants, as described in our previous work (Sun, Yu, et al., 2017a) (**Figures S1**). This subjective ambiguity model obtained the same main effect of ambiguity and category × ambiguity interaction effect on drift rate (**Figure S3d**). Although subjective ambiguity ratings deviate from the morphing-defined ambiguity levels (**Figure S1**), both modeling converge on highly similar drift-rate patterns. This consistency demonstrates that our key findings are robust regardless of whether ambiguity is defined objectively by morph levels or subjectively by participants’ perceptual judgments, confirming that increased ambiguity reliably slows evidence accumulation. Together, these findings indicate that as the ambiguity level increases, the evidence accumulation slows down, highlighting the potential of the drift rate to capture the dynamics of the decision-making process under ambiguity.

We next examined the DDM results based on the winning neural model. As expected, in **Experiment 1** (**Figure 2c**), our results revealed a significant positive interaction between category and LPP in the facial expression judgment task (99.9%). The effect size of LPP × category interaction was significantly larger and more positive in the facial expression judgment task than in the other two tasks (99.9%), highlighting the robustness of ambiguity sensitivity when the stimulus-response mapping is congruent. Although previous work has shown that LPP magnitudes were comparable between the facial expression judgment task and the wealth judgment task (Sun, Yu, et al., 2017a) (**Figures S2c, d**), the LPP-informed drift diffusion model dissociated the two tasks, indicating that DDM offers a more sensitive characterization of LPP’s role in the hidden cognitive processes underlying different types of decision-making under perceptual ambiguity. Notably, the significant positive LPP × category interaction was replicated in **Experiment 2a**, where the same fearful-happy morphed face stimuli and facial expression judgment task were combined with non-speeded responses (**Figure S3e**). These results provide support for the hypothesized role of the LPP in social-affective decision-making under perceptual ambiguity and demonstrate that the LPP tracks perceptual ambiguity primarily when the ambiguity is relevant to the current decision-making task.

To ensure that our central conclusions were not driven by potential effects on decision thresholds, we conducted additional model comparisons allowing the boundary parameter to vary as a function of stimulus category, ambiguity level, or their interaction. Models in which only the threshold—rather than the drift rate—was modulated by experimental manipulations consistently fit the data worse than drift rate-modulated models (**Table S3**). Moreover, adding ambiguity-dependent threshold term to the best-fitting drift-rate model, while making the model more complex, did not improve model performance. Together, these results suggest that ambiguity-related changes in decision-making processes are best captured by modulation of the drift rate, rather than by shifts in decision boundary.

### Domain-general and context-sensitive evidence accumulation

We next examined whether the role of the LPP in social-affective decision-making under perceptual uncertainty can be generalized to other types of categorization decisions under perceptual ambiguity. In **Experiment 2b**, participants viewed angry-disgust morphed face images and made angry/disgust binary choices. In **Experiment 2c**, participants viewed cat-dog morphed images and made cat/dog binary choices. Here, RTs varied as a function of stimulus ambiguity levels in the same way as in the above experiments (Sun, Yu, et al., 2017a; Sun, Zhen, et al., 2017). We applied the above winning behavioral and neural models to the data in these two experiments and found the same patterns as in **Experiment 1a**: both the category `×` ambiguity modulation effect and the category `×` LPP modulation effect on drift rate were significant (posterior probabilities > 95%; **Figure 3b, 3c**). These results suggest that the role of the LPP in decision-making under perceptual ambiguity is not specific to one type of stimulus.

As an exploratory analysis, we compared the interaction effect size across the three tasks. Specifically, for the behavioral model, the probability that the category-by-ambiguity modulation effect on drift rate differed across tasks was not significant (**Figure 3b**; the difference between the fearful-happy task and the angry-disgust task: 91%; the difference between the fearful-happy task and the cat-dog task: 87%; the difference between the angry-disgust task and the cat-dog task: 56%). For the neural model, the probability that the category-by-LPP modulation effect on drift rate differed across tasks was not significant for the comparison between the angry-disgust task and the cat-dog task (**Figure 3c**; 76%) or between the fearful-happy task and the angry-disgust task (88%). The comparison between the fearful-happy and the cat-dog task in terms of the category-by-LPP modulation effect was significant (98%).

We then tested whether and how the evidence accumulation process would be modulated by varying contexts involving ambiguity and those without ambiguity (**Experiments 3** and **4**; **Figures 4a, 4b**). Our previous work indicates that when perceptual ambiguity is no longer relevant for the participants’ decision-making, LPP does not track the perceptual ambiguity of the stimuli *per se*. For example, in a task where angry-disgust morphed faces and cat-dog morphed animals were presented, and the participants’ task was to judge whether a presented image was a human face or an animal, LPP magnitude did not track with the level of perceptual ambiguity within each domain (e.g., percentage of anger in the angry-disgust morphing face, or percentage of cat in the cat-dog morphing image; **Experiment 3**; **Figure 4a**). This is understandable because in this task, within–domain perceptual ambiguity is not relevant to the decision-making task – no matter how ambiguous a face is in terms of facial expression, it is clearly a human, not an animal. Here, we examined whether the model-based indicator of the evidence accumulation process, namely, the modulation effect on drift rate, reflects this context-dependent nature of LPP in perceptual decision-making. As in the above experiments, for the behavioral model, we included within–domain perceptual ambiguity level (i.e., fearful-happy morph level and cat-dog morph level), within–domain category (i.e., fearful vs. happy, and cat vs. dog), and their interaction as modulating factors of drift rate. For the neural model, we replaced the within–domain perceptual ambiguity level with LPP magnitude. Since the decision was at the between-domain level (i.e., judging human vs. animal), we predicted that the within-domain perceptual ambiguity and the within-domain category should not matter for the decision. As predicted, we did not observe significant interaction effect in neither the behavioral model nor the neural model (posterior probability of the interaction effect smaller than 0: within–domain category × ambiguity = 35%, within–domain category × LPP = 26%; **Figures 4c, d**).

To further examine the context-dependent nature of LPP’s role in resolving perceptual ambiguity, we re-analyzed another dataset using DDM (**Experiment 4**). In this task, participants viewed facial images expressing happy or fearful emotions in three consecutive blocks (Block 1 through Block 3). The middle block (Block 2) was the same as Experiment 2, where the participants viewed fearful-happy morphed faces and judged whether a presented image depicts fearful or happy expression. In Block 1 and Block 3, only unambiguous fearful and happy facial images were presented. Our previous work showed that LPP elicited by the unambiguous stimuli was reduced in the perceptually certain context (i.e., Block 1 and Block 3) relative to the perceptually ambiguous context (i.e., Block 2), again suggesting the context-dependent nature of LPP’s role in resolving perceptual ambiguity) (Sun, Zhen, et al., 2017). Here, we predicted that the category `×` LPP modulation on drift rate should be larger in the perceptually ambiguous context than in the perceptually unambiguous context. To test this hypothesis, we first included only the unambiguous facial images (100% fearful or happy) in each category, so that we could compare the LPP-by-category modulation effect within the same stimuli across all three blocks. As we predicted (**Figures 4e**, **4f**), the category `×` LPP modulation on drift rate was significantly higher in the perceptually ambiguous context (Block 2) than in the first perceptually unambiguous context (Block 1; posterior probability > 99%). Interestingly, we observed an increasing trend in the category `×` LPP modulation on drift rate in Block 3 relative to Block 1 (posterior probability = 94.8%), possibly reflecting a carry-over effect from Block 2. For completeness, we found that the category `×` LPP modulation on drift rate across all levels showed similar results to those in Experiment 2a (**Figure 4e)**. Altogether, these results suggest that the role of LPP as the neural mechanism of resolving perceptual ambiguity in social-affective decision-making is context dependent – LPP does not track perceptual ambiguity *per se* but does so only when resolving perceptual ambiguity is relevant to the present task.

Notably, for completeness, we also reported the traditional behavioral results, including RTs, confidence rating, and response accuracy across ambiguity levels and stimulus categories in the *Supplementary Materials* (see **Figure S5** through **Figure S9** for more details). The grand average ERPs by ambiguity levels across experiments (**Figure S6**), the comparison with the Pz as a main electrode and the centro-parietal as a cluster (**Figure S7**), together with scalp topographies of ambiguity-related ERP differences in the speeded and non-speeded emotion judgment task (**Figure S8**).

### Single neuron activities from the amygdala and dmPFC reflect evidence accumulation

In **Experiments 1-4**, we show that the ERP component LPP reflects the neurocognitive processes underlying ambiguity resolution in social-affective decision-making under perceptual ambiguity. Previous research has suggested cortical origins of the LPP (Sun, Zhen, et al., 2017). However, source localization based on EEG signals is challenging and inconclusive. Single-unit recordings address these limitations by directly measuring neuronal electrical activities from single neurons. Therefore, in **Experiment 5**, we aimed to provide a finer-grained mechanistic understanding of the neuronal basis of the decision-making process by combining single-unit recordings and DDM. In particular, we recorded single-neuron activities in a group of epilepsy patients while they performed the same task as in **Experiment 2** (see **Table S1** for demographic details). Specifically, we recorded neuronal activities from two brain regions consistently involved in decision-making and processing emotional information – the bilateral amygdala and dmPFC (**Figure 5a**).

Previous research has consistently implicated the amygdala in encoding both the emotional valence of facial expressions and perceptual ambiguity (Adolphs et al., 1994, 1999). Specifically, using the same fear-happy morphed face stimuli, a previous single-unit recording study has shown that amygdala neuronal activities track the decreasing ambiguity level of facial expression (S. Wang et al., 2017). We therefore hypothesized that, similar to the role of the LPP in **Experiments 1-4**, here trial-by-trial neuronal activities in the amygdala should have a significant positive interactive effect with stimulus category on drift rate. Similarly, previous work has shown that neuronal activities in dmPFC are sensitive to increasing decision ambiguity and uncertainty (Alexander & Brown, 2010; De Martino et al., 2012; Rushworth & Behrens, 2008; Shackman et al., 2011; Sheth et al., 2012), and this brain region has been consistently identified as the source of the LPP (Sun, Yu, et al., 2017b; Sun, Zhen, et al., 2017). Given the opposing patterns in ambiguity processing in dmPFC neurons in comparison with amygdala neurons (**Figures 5b**, **5c**, **5d**, **5e**), we hypothesize that their encoding of drift rates will also differ correspondingly. Specifically, we predicted that dmPFC neurons would exhibit a negative interaction effect with stimulus category on drift rate, and vice versa for amygdala neurons.

The behavioral DDM replicated the results observed in the above EEG experiments. Specifically, the interaction effect of ambiguity level and stimulus category on drift rate was significantly lower than 0 (probability > 99%). This result confirmed that the drift rate in our DDM model remained a good indicator of the evidence accumulation process in decision-making under ambiguity in this group of neurosurgical patients.

We then proceeded to examine the role of amygdala and dmPFC neuronal activities in resolving ambiguity in decision-making. The model with amygdala activity revealed a significant and positive category `×` firing rate interaction (**Figure 5f**, probability > 99%). In contrast, the category `×` firing rate interaction effect on drift rate was significantly negative for dmPFC neurons (**Figure 5g**, probability > 99%). These results support our hypothesis and collectively indicate the roles of the amygdala and dmPFC in encoding decision parameters and highlight the functional dissociation of both regions at the single-neuron level.

### Functional connectivity between the amygdala and dmPFC correlates with individual drift rate

**Experiment 5** demonstrated that single neurons in the dmPFC and amygdala reflected evidence accumulation processes in social-affective decision-making under perceptual ambiguity. **Experiment 6** aims to examine the neural correlates of decision parameters within the DDM framework at the network level using fMRI and to further elucidate how these parameters are encoded by functional connectivity between the amygdala and dmPFC. Previous work has shown that the right amygdala is functionally connected to the dmPFC, and their connectivity strength is associated with ambiguity resolution in perceptual decision-making (Sun, Yu, et al., 2023). Here, we investigated whether the PFC-amygdala connectivity strength may explain individual differences in the computational index of the evidence accumulation process (i.e., drift rate in the DDM model). Participants (N = 19) completed the same fearful-happy facial expression judgment task as in **Experiment 2** in an MRI scanner.

We first applied the winning behavioral DDM model to the behavioral data of the fMRI study. Consistently, we observed a negative category `×` ambiguity interaction effect on drift rate among the fMRI participants (**Figure 6a**). To further examine whether and how the drift rate was encoded by the frontal-amygdala circuit, we extracted individual category `×` ambiguity interaction effects from fMRI participants and tested their correlation with their right amygdala-dmPFC connectivity.

**Figure 6.**
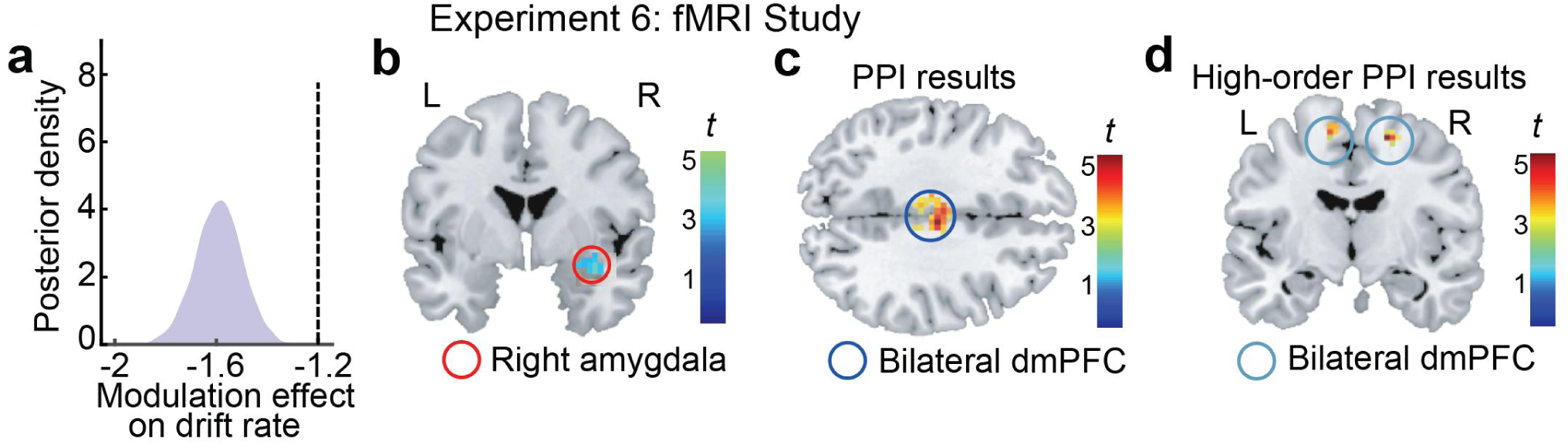
Results of functional connectivity analysis in the fMRI study (Experiment 6). **(a)** Behavioral drift rates in fMRI participants. **(b)** The right amygdala was involved in ambiguity processing. **(c)** Bilateral dmPFC/pre-SMA regions were co-activated with the right amygdala during ambiguity processing. **(d)** Bilateral dmPFC/pre-SMA regions were co-activated with the right amygdala, and their connectivity strength was correlated with individual differences in the modulation effect on drift rate.

We first performed a psychophysiological interaction (PPI) analysis with ambiguity levels as the psychological factor to identify target regions functionally connected to the seed region, right amygdala (**Figure 6b**; MNI peak: *x* = 30, *y* = 0, *z* = –21), using signals from a 6-mm-radius sphere around the seed as the volume of interest. The peak coordination was reported in our previous work, which showed decreasing activity with increasing levels of ambiguity (Sun, Yu, et al., 2017a; Sun, Zhen, et al., 2017; Sun, Yu, et al., 2023). We then examined whether this connectivity was correlated with individual category `×` ambiguity interaction effects on drift rates. Whole-brain PPI analysis revealed that the right amygdala showed co-activation with the bilateral dmPFC (**Figure 6c**; MNI peak: *x* = 0, *y* = –15, *z* = 39; 14 voxels, SVC, *P*FWE = 0.014) under conditions of a positive PPI interaction. However, no significant correlation was observed between PPI connectivity strength and individual drift rates.

We then conducted a higher-order PPI analysis to identify brain regions functionally connected to the right amygdala, whose connectivity was modulated by individual category `×` ambiguity interaction effects on drift rates derived from the DDM. Specifically, the individual interaction effect on drift rate was used as a second level (i.e., participant level) regressor to identify regions responsive to the primary PPI contrast. This analysis revealed co-activation of the seed region (right amygdala) with the right dmPFC/pre-supplementary motor area (pre-SMA) (**Figures 6d**; MNI peak: *x* = 18, *y* = –9, *z* = 57; Z = 4.12; 16 voxels). A non-parametric Spearman’s correlation analysis further confirmed a strong association between right amygdala–right dmPFC connectivity strength and individual drift rates, with stronger connectivity associated with a stronger negative interaction effect on drift rates. While the voxel-wise correlation implemented in SPM relies on Pearson statistics, which are sensitive to distributional assumptions and outliers, the Spearman’s approach provides a more distribution-free estimate. Additionally, we identified activation in the left dmPFC/pre-SMA (**Figures 6d**; MNI peak: *x* = –12, *y* = –9, *z* = 60; Z = 3.57; 10 voxels), which was also co-activated with the right amygdala.

Together, these findings highlight how the frontal–amygdala network contributes to decision-making under ambiguity, as indexed by the DDM-derived decision parameters. Notably, previous work has shown that the dorsal medial frontal cortex (dmMFC)—which encompasses both the anterior and posterior mid-cingulate cortex (MCC) and overlaps with the pre-supplementary motor area (pre-SMA)(Clairis & Lopez-Persem, 2023; de la Vega et al., 2016; Dixon et al., 2017; Vogt et al., 2003)—plays a critical role in self-monitoring and the control of actions (Fu et al., 2019; Ullsperger et al., 2014; Vogt et al., 2003). In line with this established functional and anatomical organization, our direct comparison of intracranial contact locations with fMRI activations demonstrates that, although the fMRI cluster is slightly more posterior and extends toward the posterior MCC or pre-SMA (**Figure S10**), both sets of sites fall within the broader dmMFC.

## Discussion

Our study used drift diffusion modeling (DDM) to integrate behavioral, EEG, single-neuron, and fMRI connectivity findings on perceptual decision-making under emotional ambiguity. Across tasks, increasing ambiguity was associated with lower drift rate, indicating less efficient evidence accumulation. This effect was strongest when ambiguity was relevant to the judgment, weaker when emotional cues indirectly informed wealth judgments, and absent or substantially reduced when ambiguity was orthogonal to the task. Model comparisons further showed that ambiguity-dependent behavior was better captured by drift-rate modulation than by boundary shifts, suggesting that perceptual ambiguity primarily affected the quality of accumulated evidence rather than decision caution. Analysis of EEG and single-unit recording data links this parameter to specific neural signatures, including the LPP and single-neuron activity in the dmPFC and amygdala, revealing the neural substrates underlying the evidence accumulation process involved in perceptual decision-making under ambiguity. Together, these findings support the view that evidence accumulation is dynamically regulated by the dmPFC— and potentially the broader PFC-amygdala circuit — highlighting the role of the corticolimbic system in social-affective perceptual decision-making and extending beyond existing theoretical frameworks.

A noteworthy point is that parts of the neural datasets were reported in previous studies. Those studies showed that the late positive potential (LPP) tracks emotional ambiguity (Sun, Zhen, et al., 2017), that amygdala and medial frontal neurons respond to ambiguity (S. Wang et al., 2017), and that amygdala–medial frontal connectivity is related to ambiguity processing (Sun, Yu, et al., 2023). The present study builds on these findings in a specific way. Rather than treating these neural effects as separate markers of ambiguity, we asked whether they can be understood in terms of a common computational framework derived from behavior. In this sense, the new contribution of the current work is not the discovery of ambiguity-sensitive neural activity *per se*, but the linking of these neural responses to evidence accumulation as measured by DDM.

This framing is consistent with a growing literature applying evidence accumulation models to social-affective decision-making. Previous work has shown that emotional intensity and ambiguity in face categorization influence drift rate and decision bias (Haller et al., 2024). Yau and colleagues showed that facial emotion information decoded from the fusiform gyrus modulates drift rate during dynamic happy–sad categorization, and that centroparietal positivity tracks evidence- and urgency-related signals in a face-morphing task (Yau et al., 2020, 2021). El Zein et al. (2024) showed that threat-related facial information can be processed earlier than a matched non-emotional dimension (Zein et al., 2024). Our findings therefore do not suggest that DDM is new to emotion perception. Rather, they show that DDM can be used to bring together EEG, single-neuron, and fMRI connectivity results that were previously described mainly in terms of ambiguity.

The EEG results are informative in this regard. In affective neuroscience, the LPP has often been associated with sustained attention to emotionally salient stimuli (Hajcak et al., 2010, 2011; Moratti et al., 2011; Schupp & Kirmse, 2022). In perceptual decision-making, a similar centroparietal positivity has been described as a neural signal of evidence accumulation (Kelly & O’Connell, 2013; R. G. O’connell et al., 2012; Yau et al., 2021). Our results link these two traditions. Trial-by-trial LPP amplitude was associated with drift rate, but this relationship depended on task-relevance of ambiguity. The association was strongest when participants judged emotional expression, and much weaker when the same facial ambiguity was irrelevant to the decision. This pattern argues against a simple interpretation that the LPP only reflects visual ambiguity or emotional salience. Instead, the LPP appears to reflect the extent to which emotional information is used as decision evidence.

The single-neuron findings further suggest that amygdala and medial frontal activity are related to different aspects of ambiguity processing. Amygdala firing was positively associated with drift rate. This is consistent with the idea that the amygdala is sensitive to the availability of emotion-relevant evidence, especially for threat-related or affectively salient facial cues (M. J. Kim et al., 2022; Méndez-Bértolo et al., 2016; Y. Wang et al., 2023). In contrast, medial frontal firing showed a negative association with drift rate. This pattern is compatible with the role of medial frontal regions in uncertainty, conflict, and monitoring when available evidence is weak. Thus, the two regions did not simply show the same ambiguity effect in different directions of measurement. Rather, their activity may index complementary aspects of the decision: the amygdala tracking category-diagnostic affective evidence, and medial frontal cortex tracking the difficulty or conflict generated by weak evidence.

The fMRI connectivity result should be interpreted with caution. Stronger amygdala–medial frontal coupling was associated with stronger ambiguity-related modulation of drift rate. This suggests that individual differences in interactions between affective and medial frontal systems are related to how efficiently ambiguous emotional evidence is accumulated. At the same time, the medial frontal regions identified across the single-neuron and fMRI datasets are not anatomically identical. Some fMRI clusters extend toward medial frontal and premotor regions, which have been linked to action selection, urgency, and competition between response options (Cisek, 2007; Cisek & Kalaska, 2010; Thura et al., 2025; Yau et al., 2020). We therefore interpret the fMRI findings as evidence for a broader amygdala–medial frontal/premotor network associated with decision formation, rather than as strict anatomical convergence on one dmPFC location.

Several limitations should be noted. First, many analyses were performed on previously collected datasets, and the present work is primarily a computational reanalysis. Second, the neural findings are correlational. Although convergence across EEG, single-neuron activity, and fMRI connectivity strengthens the interpretation, the data do not show that amygdala or medial frontal activity causally changes drift rate. Third, the non-speeded tasks are limited because participants may have formed their decisions before making a button press. The similar drift-rate pattern in the speeded task reduces this concern, but future studies should use designs that separate evidence accumulation from motor preparation more cleanly.

Taken together, these findings suggest that emotionally ambiguous decisions can be understood as changes in evidence accumulation. The study does not identify a new decision circuit. Instead, it shows that neural responses previously linked to emotional ambiguity can be related to a common computational process: the efficiency with which affective evidence is accumulated toward a categorical judgment.

## Data availability

The preprocessed data used in Experiments 1–6 were published in Sun et al. (2023) (54) and is available at https://osf.io/26rhz/files/osfstorage.

## Code availability

The DDM model code and the corresponding data are available at https://doi.org/10.17605/OSF.IO/GK328.

## Declare of interests

The authors declare no competing interests.

## Acknowledgements

The authors have no funding sources to list for this study.

## Supplementary Materials

### Supplementary Tables

**Table S1.**
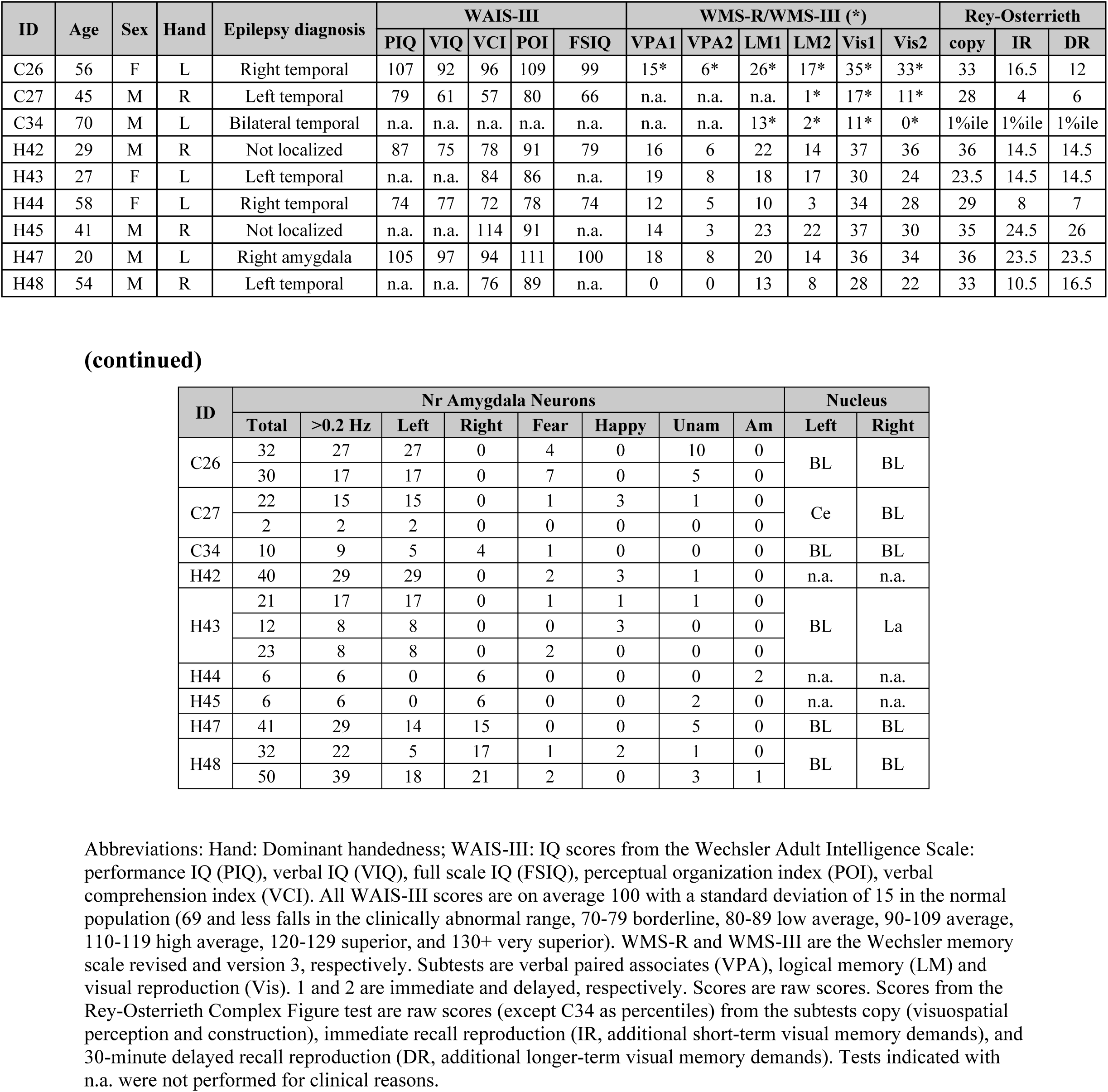
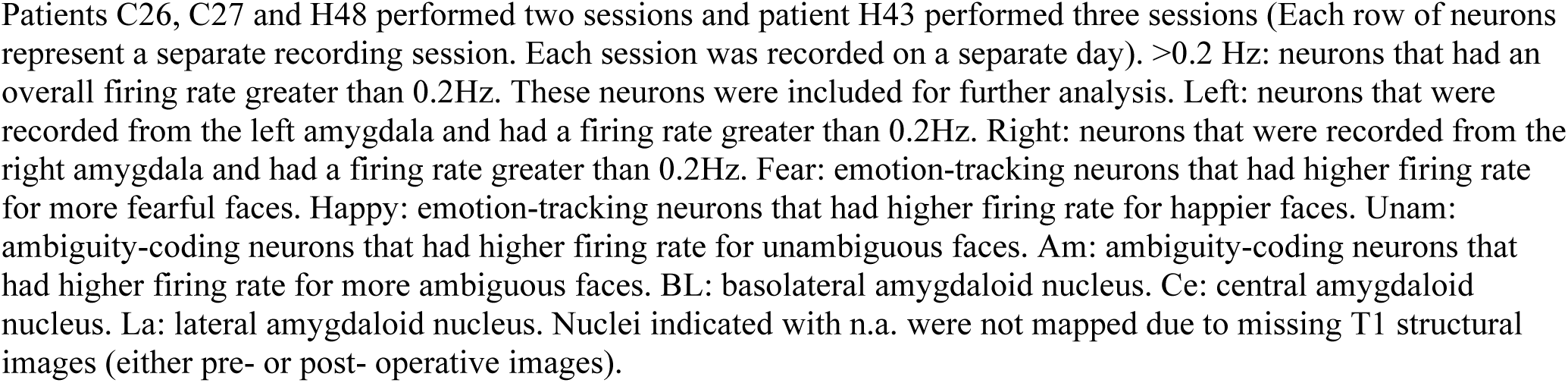
List of patient demographics, pathology, and neuropsychological evaluation (Experiment 5).

**Table S2.**
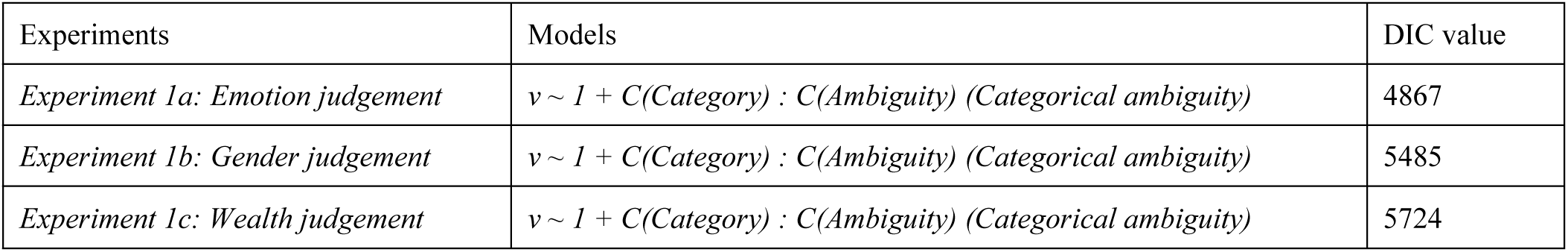
Behavioral models with separate stimulus categories and objective ambiguity levels.

| Experiments | Models | DIC value |
| --- | --- | --- |
| *Experiment 1a: Emotion judgement* | $v \sim 1 + C(\text{Category}) : C(\text{Ambiguity})$ (Categorical ambiguity) | 4867 |
| *Experiment 1b: Gender judgement* | $v \sim 1 + C(\text{Category}) : C(\text{Ambiguity})$ (Categorical ambiguity) | 5485 |
| *Experiment 1c: Wealth judgement* | $v \sim 1 + C(\text{Category}) : C(\text{Ambiguity})$ (Categorical ambiguity) | 5724 |

**Table S3.**
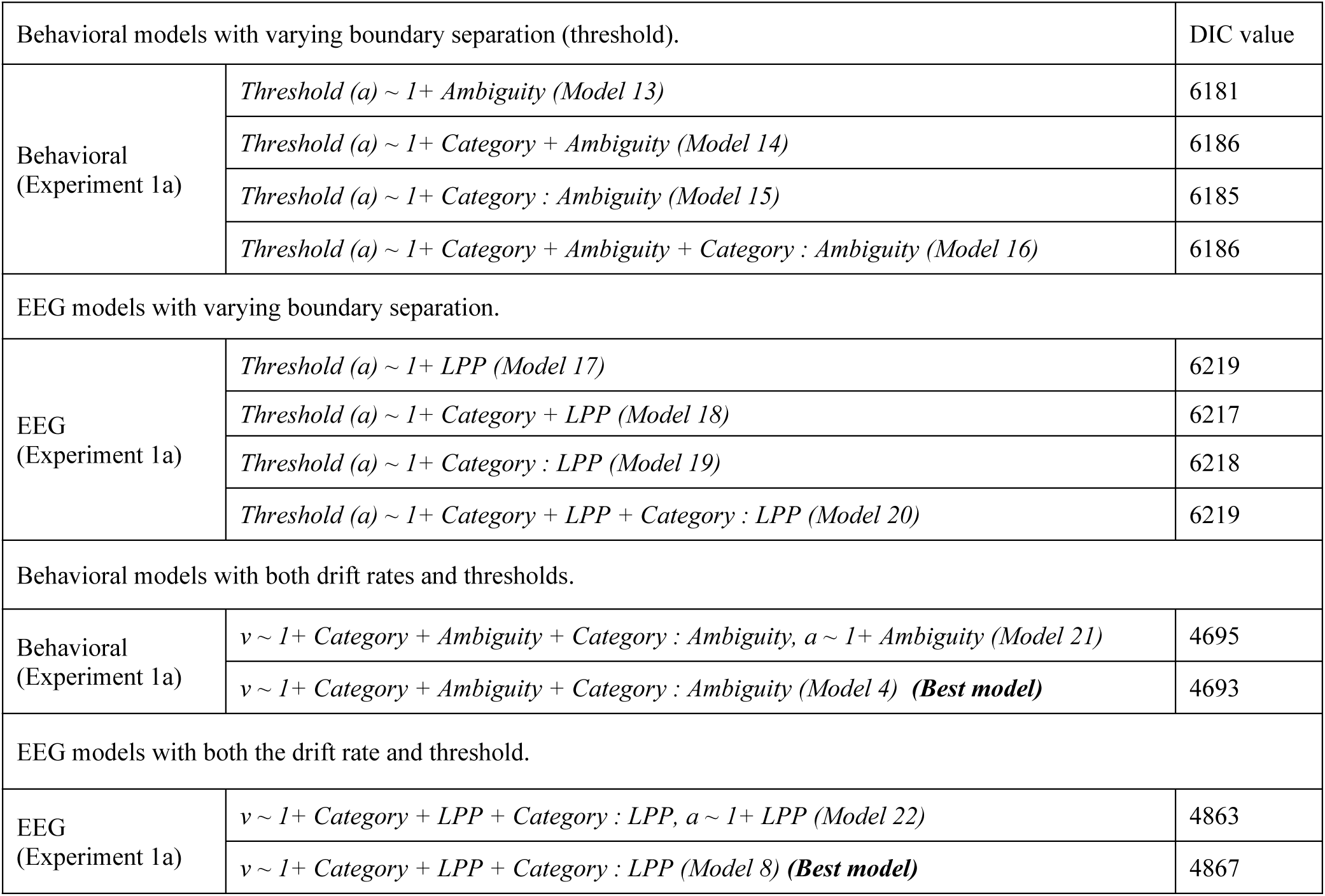
Comparisons between models with threshold modulation and models with drift rate modulation only.

### Supplementary Figures

**Figure S1.**
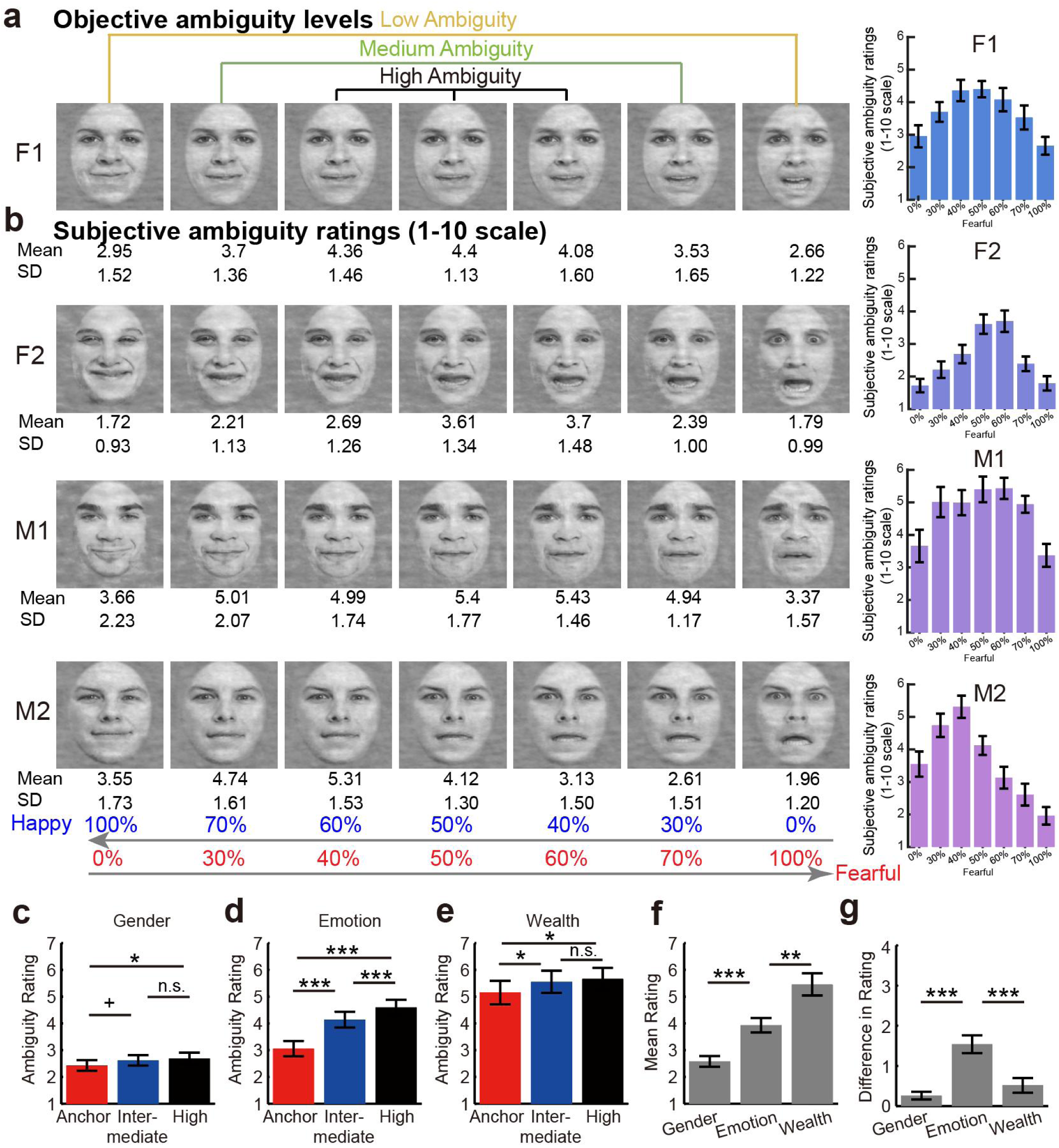
Subjective ambiguity ratings for each stimulus, across levels and tasks. (**a**) Morphed emotional face stimuli (F1, F2, M1, M2) vary from 100% happy to 100% fearful. (**b**) Subjective ambiguity ratings (1–10 scale) show a consistent inverted-U pattern across identities, with the highest perceived ambiguity at intermediate morph levels. Error bars indicate ± s.e.m. (**c–e**) Ambiguity ratings are shown for Gender, Emotion, and Wealth judgments across anchor, intermediate, and high ambiguity levels. The Wealth task received the highest overall ambiguity ratings (**e**, **f**), whereas the Gender task showed the lowest (**c**, **f**). The Emotion task exhibited the largest separation across ambiguity levels (**d**), consistent with the direct stimulus–response mapping that makes ambiguity manipulation most perceptually salient. Difference scores (**g**) confirm that ambiguity effects were strongest for Emotion, moderate for Wealth, and minimal for Gender. Error bars indicate ± s.e.m. \**p* < .05*, ** p < .01*, *** *p* < .001, ^+^*p* < .10, n.s. indicates not significant. Adapted from Wang et al., 2017, Nature Communications.

**Figure S2.**
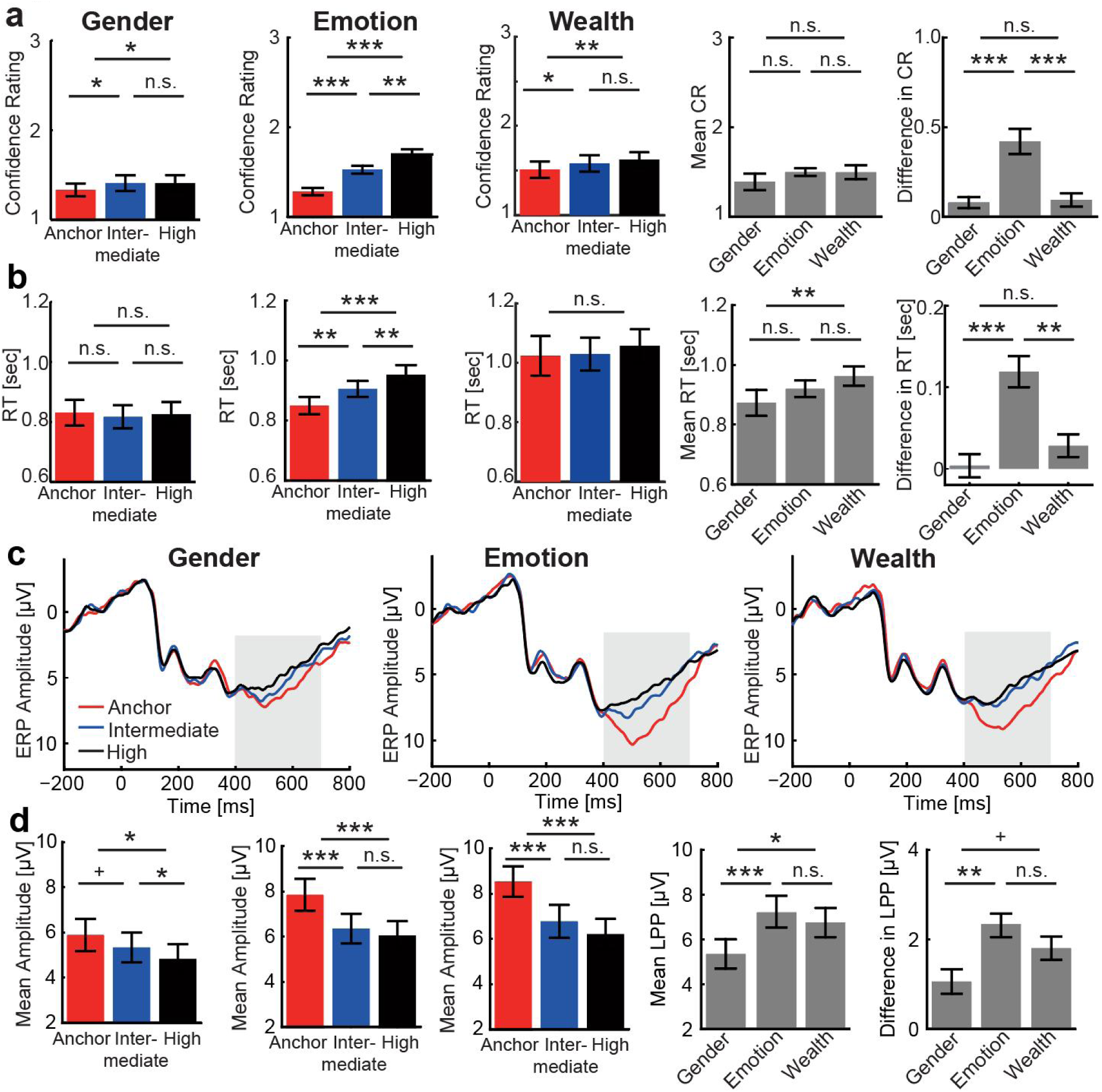
Behavioral and ERP effects across Gender, Emotion, and Wealth judgments in Experiment 1. **(a-b)** Confidence rating (CR, where 1 = most confident, 3 = least confident) and reaction time (RT) vary systematically with ambiguity level (Anchor = no ambiguity, Intermediate = moderate ambiguity, High = highest ambiguity). The strongest effects of ambiguity on confidence ratings and reaction times (high > anchor) were observed in the facial expression judgment task, where the ambiguity was relevant to the task. (**c**) ERP waveforms at Pz show corresponding modulation in the late positive potential (LPP; 400–700 ms window, shaded). (d) Bar plots summarize mean amplitudes and LPP responses, demonstrating robust ambiguity-related effects for Emotion and Wealth, and modest effect for Gender. Cross-task comparisons reflect the mean and difference in CR, RT, and LPP amplitudes across ambiguity levels. Adapted from Sun et al., 2017, NeuroImage; Sun et al., 2017, eNeuro.

**Figure S3.**
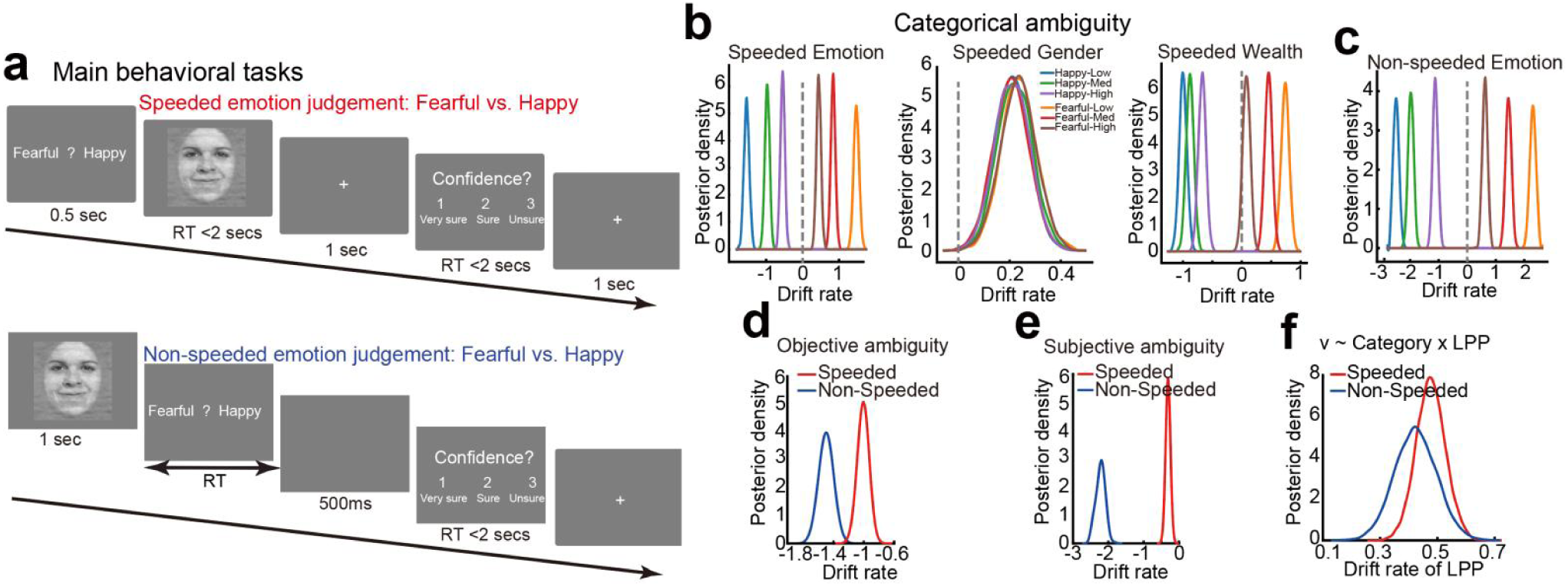
Additional task procedure and modeling results corresponding to Figure 2. (**a**) Comparison of tasks with speeded responses (Experiment 1a) and non-speeded responses (Experiment 2a). Note that the facial images shown here were adapted from a publicly available database. (**b**) Posterior drift-rate distributions by emotion category and ambiguity level across the emotion, gender, and wealth judgment tasks. Extension on Figure 2. Happy (blue, green, purple) distributions cluster on the negative side of the drift axis, whereas fearful (brown, red, orange) distributions lie on the positive side, illustrating that the sign of the interaction term reflects directional coding for both the emotion and wealth tasks. The width and separation of the distributions capture differences in evidence accumulation strength across ambiguity levels—expressed as an interaction effect—with the emotion task showing the largest separation, which becomes weaker in the wealth judgment. In contrast, the gender task shows no discernible differences in evidence accumulation across morphing levels or stimulus categories. (**c**) Posterior distribution of the category-by-objective ambiguity interaction effect for speeded (red, same as the red distribution in Figure. 2b) and non-speeded (blue, same as the red distribution in Figure. 3b) tasks. Here, objective ambiguity levels are researcher-defined. (**d**) Posterior distribution of the category-by-subjective ambiguity interaction effect for speeded (red) and non-speeded (blue) tasks. Here, subjective ambiguity levels were obtained from an independent group of participants. (**e**) The category × LPP interaction effect on drift rate showed similar patterns for the speeded and non-speeded tasks.

**Figure S4.**
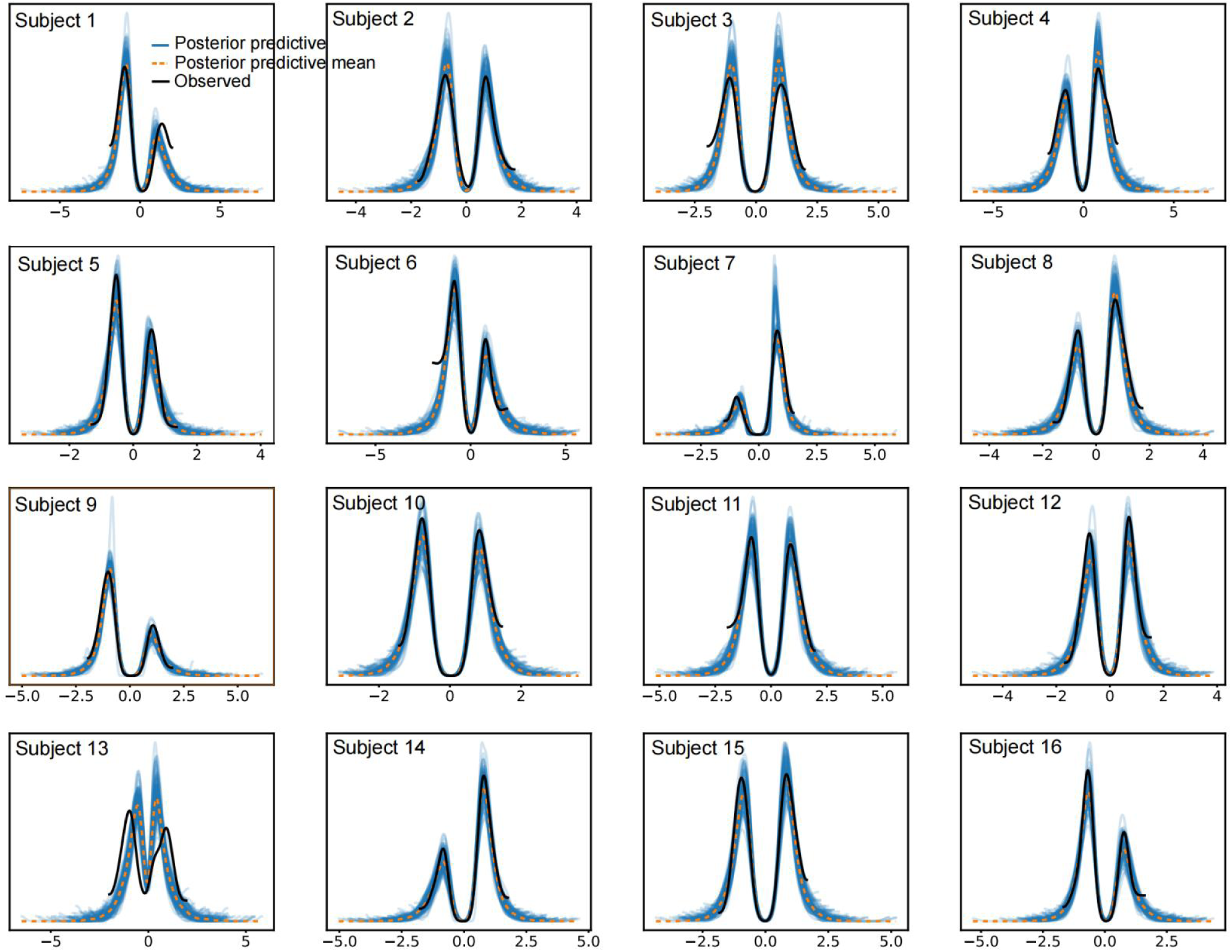
Posterior predictive reaction times (RTs) and comparison with observed RTs for each subject in the main task (Experiment 1a). Each panel represents one of the 16 individual subjects in Experiment 1a. The blue lines show posterior predictive RT distributions generated from 100 samples, which are comparable to the observed RT distributions. Overall, the model demonstrates good recovery of RT distributions, with simulated RTs closely aligning with the observed data.

**Figure S5.**
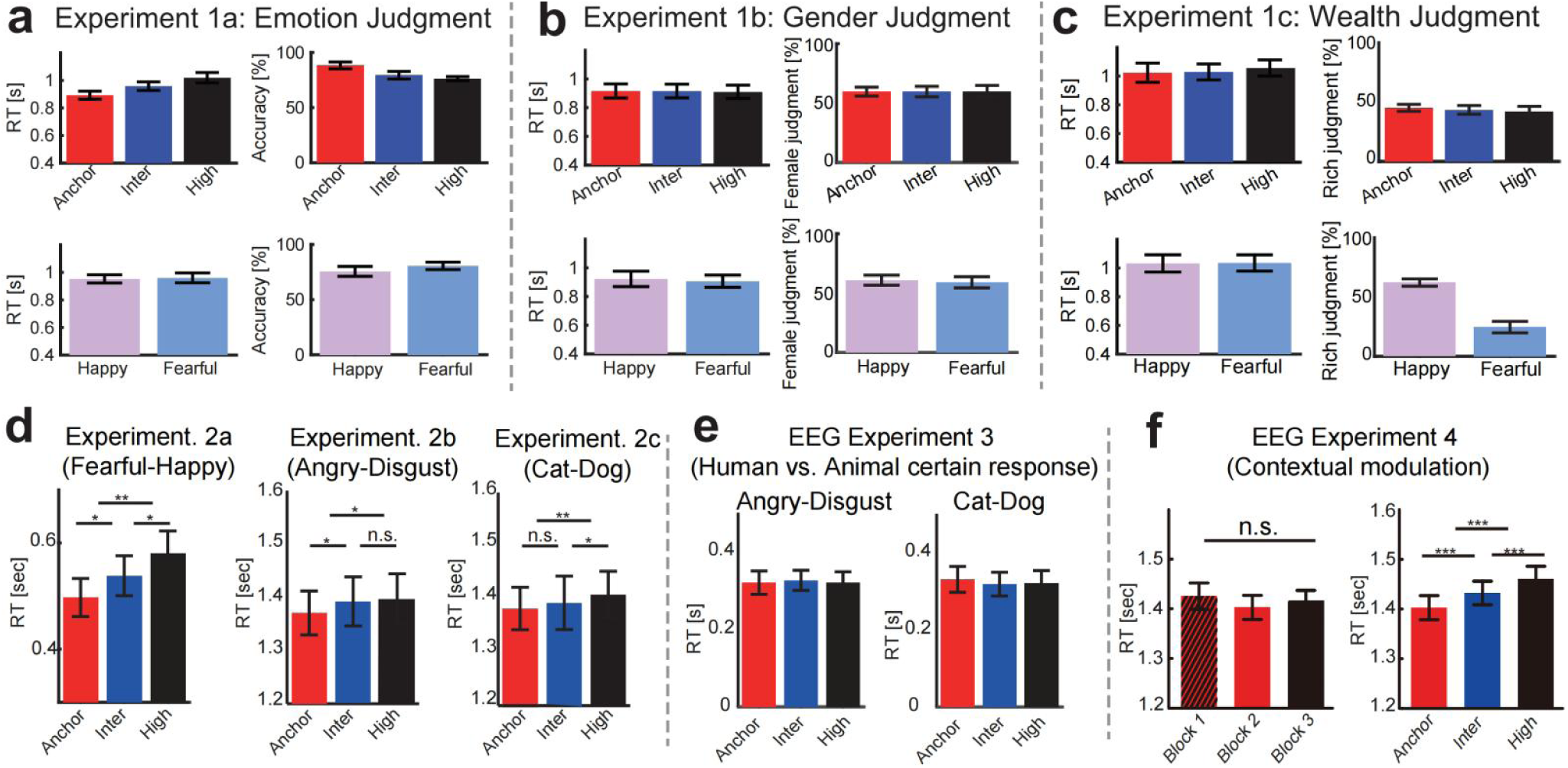
Behavioral results across Experiments 1-4. (**a-c**) RTs and behavioral responses across ambiguity levels for Experiment 1 (emotion, gender, and wealth judgments). In the emotion task (**a**), RTs increased and accuracy decreased with higher ambiguity, with no differences between fearful and happy faces. For gender (**b**) and wealth (**c**) judgments, we report the percentage of choosing one category (e.g., “female”, “rich”), which varied with ambiguity and stimulus category; happy faces were more often judged as “rich”. (**d**) In Experiments 2a-c (Fearful-Happy, Angry-Disgust, Cat-Dog), RTs increased reliably with ambiguity, showing robust ambiguity-dependent slowing across tasks with no category-specific differences. (**e**) In Experiment 3 (certain-response EEG task), RTs showed minimal modulation, consistent with the design requiring responses only when judging whether the stimulus was an animal or a human. (**f**) In Experiment 4 (contextual modulation EEG task), RTs did not differ across low-ambiguity blocks but increased systematically with ambiguity in the mixed-ambiguity block, confirming ambiguity-dependent slowing. Error bars indicate ±s.e.m. Adapted from Sun et al., 2017, NeuroImage; Sun et al., 2017, eNeuro.

**Figure S6.**
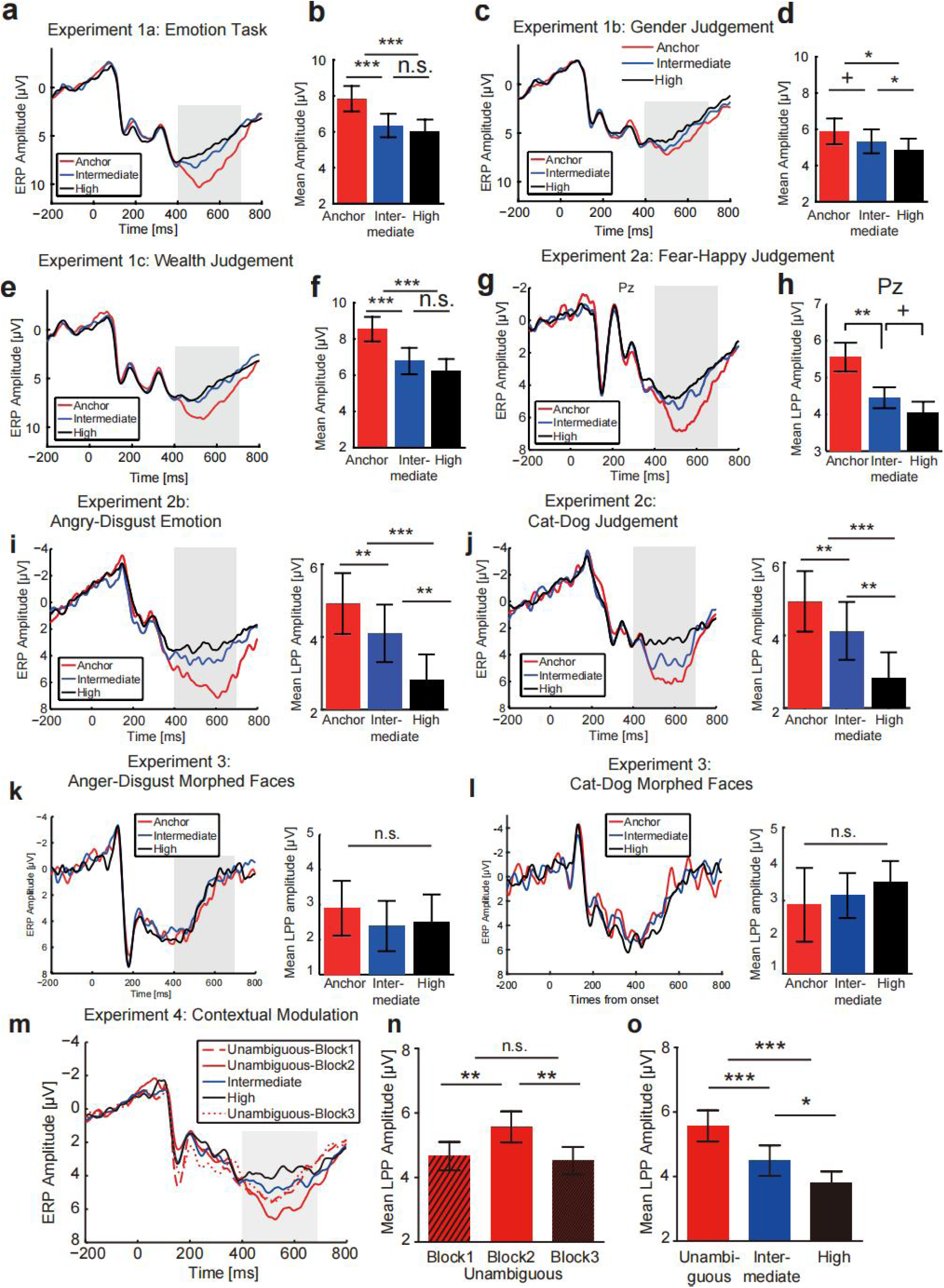
Grand-average ERPs by ambiguity levels across Experiments 1 to 4. (**a–b**) In Experiment 1a (Emotion Task), late positive potentials (LPP; 400–700 ms) increased with higher ambiguity relative to low ambiguity levels. (**c–d**) Experiment 1b (Gender Judgment) showed minimal modulation of LPP amplitude by emotional ambiguity. (**e–f**) Experiment 1c (Wealth Judgment) exhibited a similar ambiguity-dependent enhancement of LPP amplitude observed in the emotion task. (**g–h**) In Experiment 2a (Non-speeded Fearful–Happy Judgment), Pz LPP amplitude increased with ambiguity. **(i)** Experiment 2b (Angry–Disgust Judgment) showed clear LPP differences across ambiguity levels. **(j)** Experiment 2c (Cat–Dog Judgement) also showed ambiguity-related LPP modulation. (**k–l**) In Experiment 3 (morphed Angry–Disgust and Cat–Dog stimuli, with a certain response prompt), ambiguity levels did not reliably modulate LPP amplitude. (**m–o**) In Experiment 4 (contextual modulation), ERP waveforms showed no systematic block-wise differences in low ambiguous trials. However, LPP amplitude varied as a function of ambiguity levels. Across experiments, ambiguity-dependent increases in LPP amplitude were consistently observed in tasks involving emotional or socially meaningful categories, but not in decisions with low ambiguity levels. Error bars indicate ± s.e.m.; *p < .05, **p < .01, ***p < .001, n.s. indicates not significant. Adapted from Sun et al., 2017, NeuroImage; Sun et al., 2017, eNeuro.

**Figure S7.**
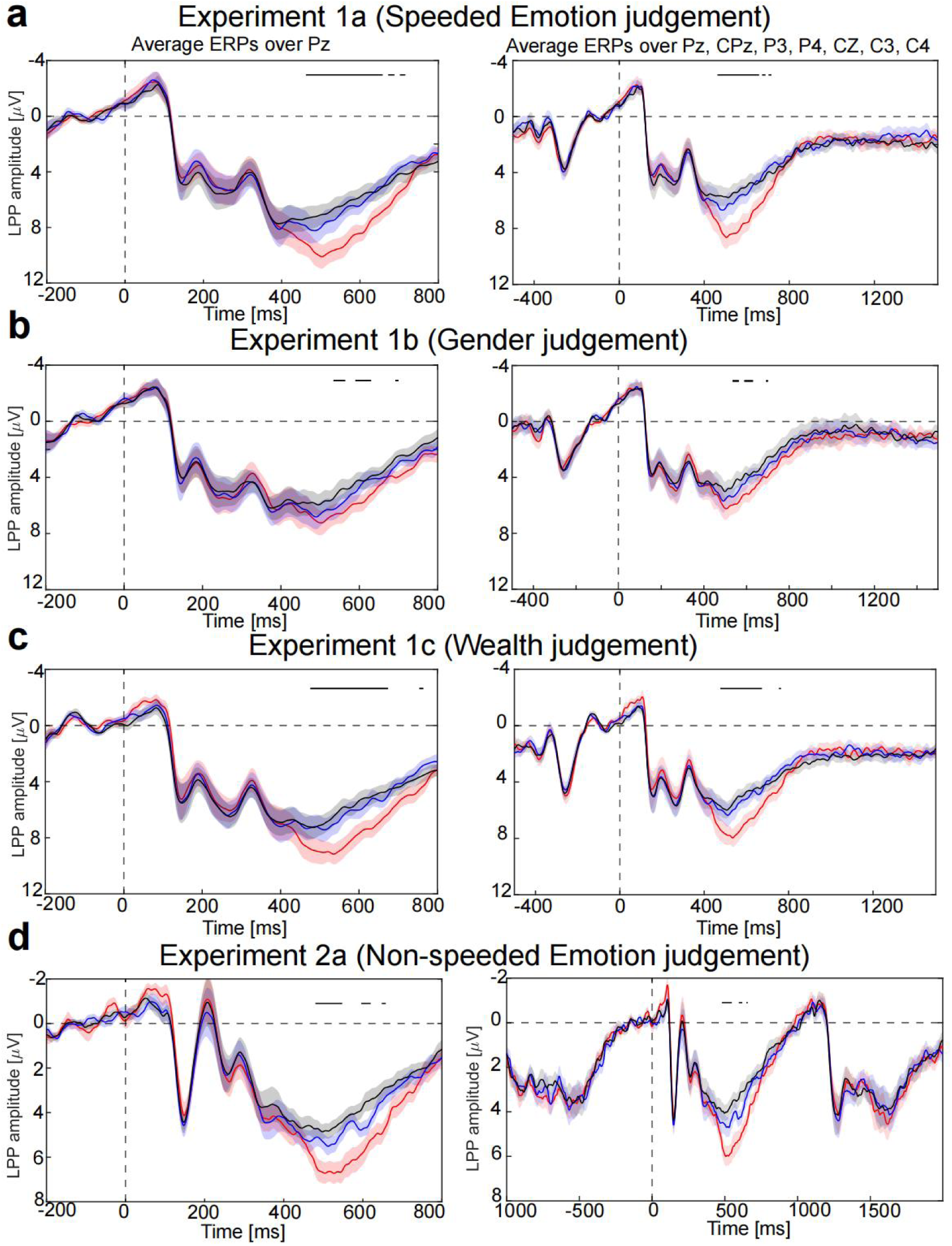
Comparison of grand-average ERP waveforms at Pz and a centro-parietal cluster. Grand-average ERP waveforms are shown for Pz (**left**) and for a centro-parietal cluster (Pz, CPz, P3, P4, Cz, C3, C4; **right**) with shaded error bars and extended windows across Experiments 1a (**a**), 1b (**b**), 1c (**c**), and 2a (**d**). The extended time window in the right panels highlights that the 400–700 ms interval consistently produces the largest ambiguity-related ERP effects. Waveforms from Pz and the centro-parietal cluster are highly similar, confirming the robustness of the ambiguity-related LPP modulation and supporting our use of the 400–700 ms mean-amplitude measure at Pz. Adapted from Sun et al., 2017, NeuroImage; Sun et al., 2017, eNeuro.

**Figure S8.**
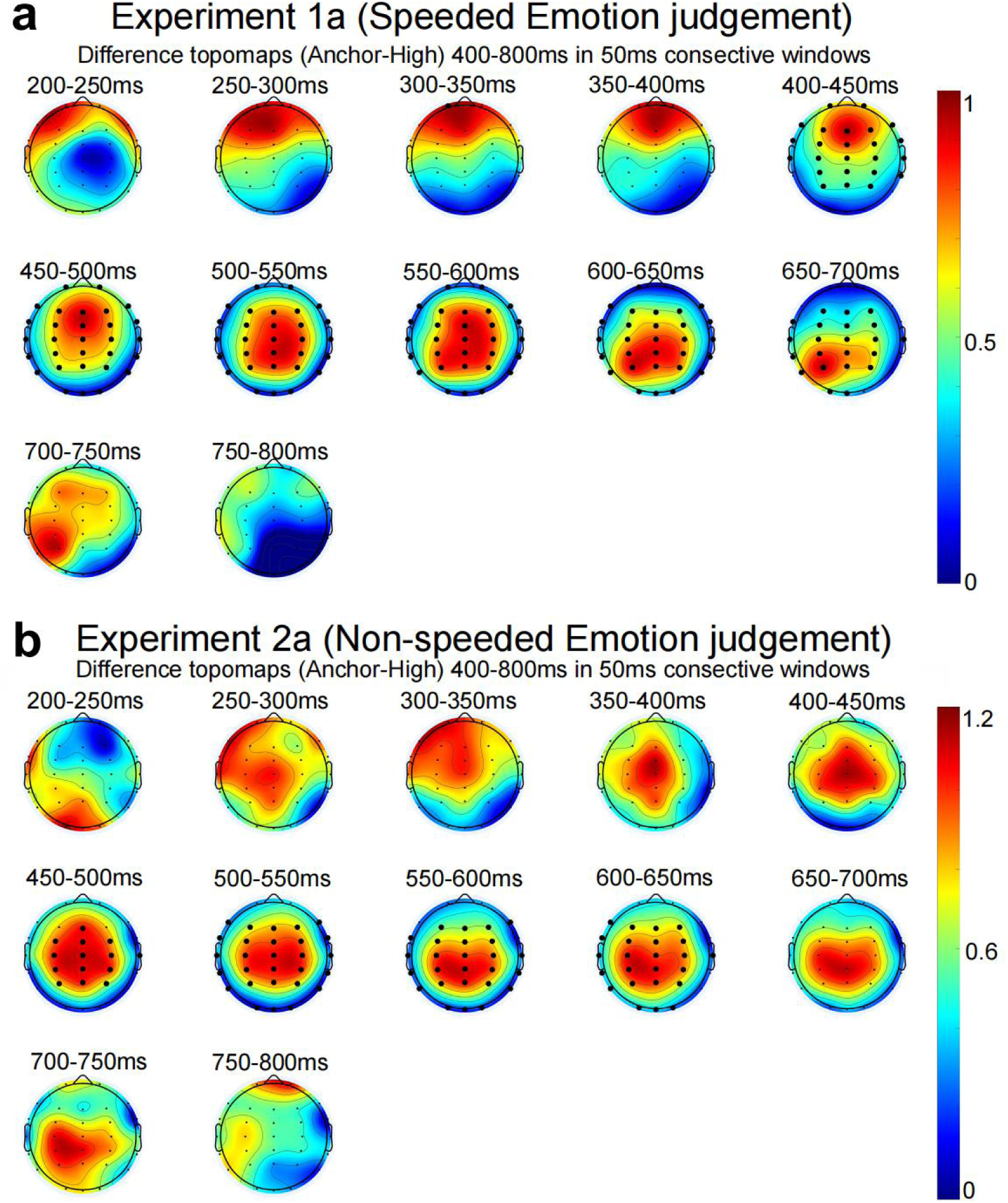
Scalp topographies of ambiguity-related ERP differences in speeded and non-speeded emotion judgments. Difference topographic maps (anchor-high ambiguity) for (**a**) Experiment 1a (Speeded emotion judgment) and (**b**) Experiment 2a (Non-speeded emotion judgment), computed across 50-ms consecutive windows from 200-800 ms after stimulus onset. Warmer colors indicate greater positivity for anchor relative to high ambiguity faces. In both experiments, robust centro-parietal positivity emerges between 400–700 ms, consistent with the chosen time window for the LPP-like ambiguity modulation. Black dots denote electrodes showing significant differences after multiple-comparison correction. Color scales reflect the difference amplitudes (anchor-high ambiguity) for each experiment. Adapted from Sun et al., 2017, NeuroImage; Sun et al., 2017, eNeuro.

**Figure S9.**
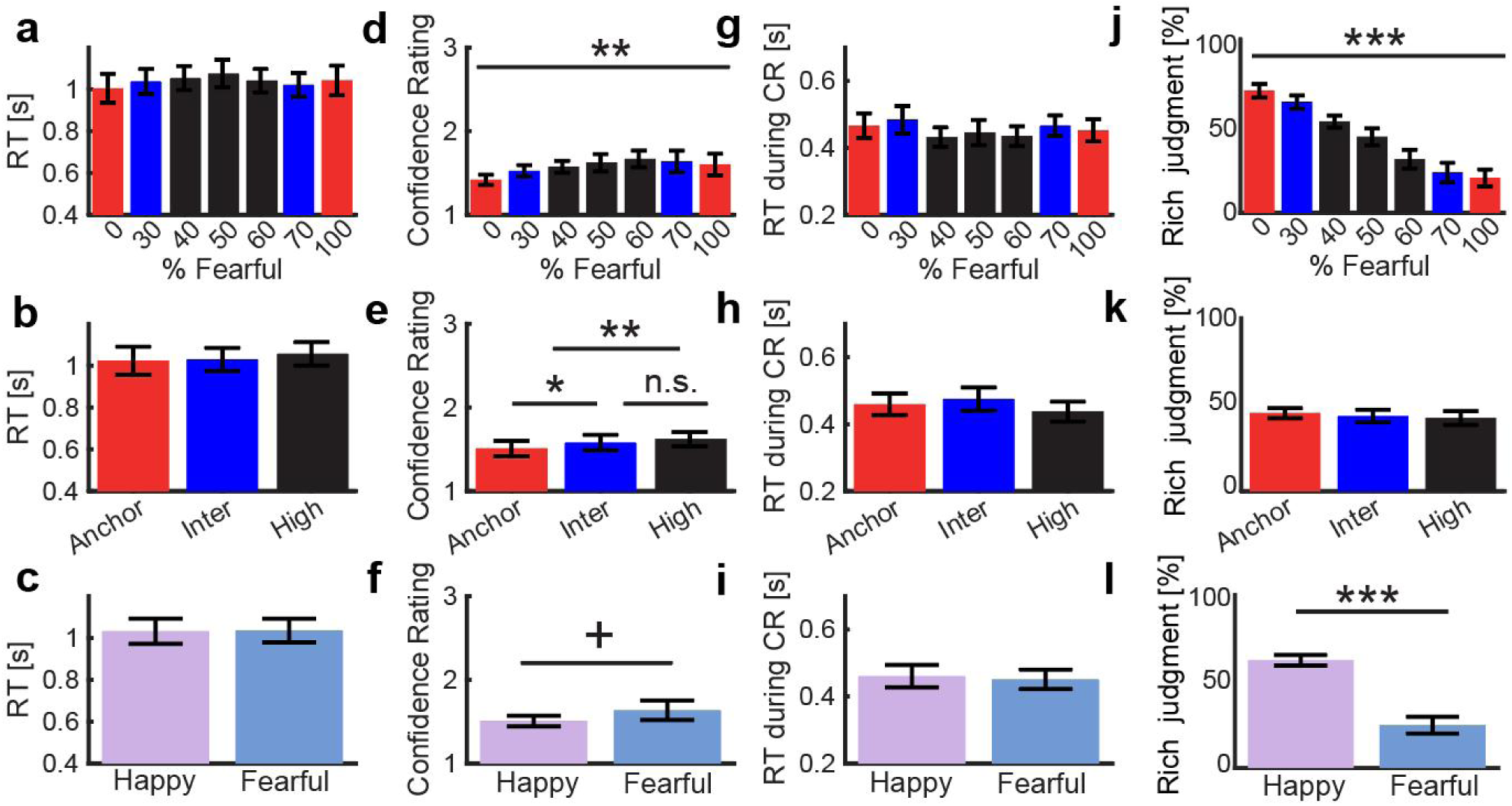
Behavioral effects of emotional expressions on wealth judgments. (**a-c**) Reaction times (RTs) showed no significant modulation by emotional ambiguity or by happy versus fearful expressions. (**d-f**) Confidence ratings increased with clearer emotional expressions: intermediate and high-ambiguity faces were rated with higher confidence than anchor faces (**e**), and clearly happy expressions elicited marginally higher confidence than clearly fearful expressions (**f**). (**g-i**) RTs during CRs showed no significant modulation by emotional ambiguity or by happy versus fearful expressions. (**j**) Happier faces tended to be judged as wealthier, whereas more fearful faces tended to be judged as poorer. (**k**) Wealth judgments were unaffected by the ambiguity level. (**l**) Wealth judgments were significantly higher for happy expressions than for fearful ones. Adapted from Sun et al., 2017, NeuroImage; Sun et al., 2017, eNeuro.

**Figure S10.**
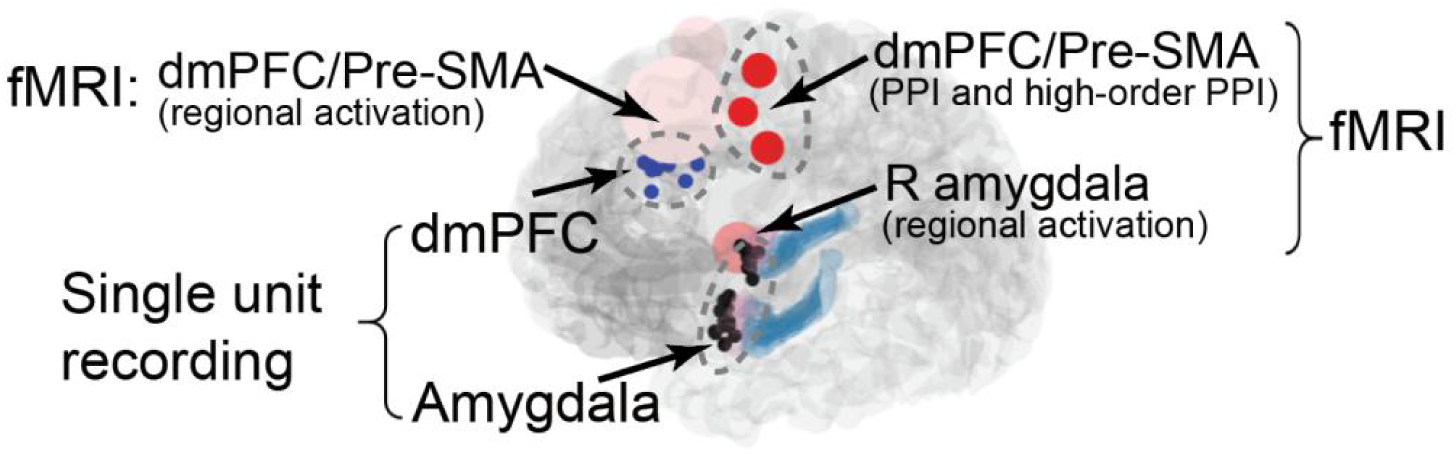
Comparison of single-unit recording sites, fMRI activation clusters. Intracranial dmPFC (blue) and amygdala (black) recording sites overlaid with fMRI activation clusters (very light red: dmPFC fMRI regional activation for decreasing ambiguity; light red: right amygdala fMRI regional activation for increasing ambiguity; dark red: dmPFC/pre-SMA activation in the PPI and higher-order PPI analyses using the right amygdala as the seed). Although the fMRI dmPFC cluster extends slightly superiorly and posteriorly toward the pre-SMA, both sets of sites lie within the broader medial frontal cortex

