## Supplementary materials for "Neural substrates of evidence accumulation in social-affective decision-making under perceptual ambiguity"

### Supplementary Tables

### Table S1. List of patient demographics, pathology, and neuropsychological evaluation (Experiment 5).

| **ID** | **Age** | **Sex** | **Hand** | **Epilepsy diagnosis** | **WAIS-III** | | | | | **WMS-R/WMS-III (*)** | | | | | | **Rey-Osterrieth** | | |
| --- | --- | --- | --- | --- | --- | --- | --- | --- | --- | --- | --- | --- | --- | --- | --- | --- | --- | --- |
|  |  |  |  |  | **PIQ** | **VIQ** | **VCI** | **POI** | **FSIQ** | **VPA1** | **VPA2** | **LM1** | **LM2** | **Vis1** | **Vis2** | **copy** | **IR** | **DR** |
| C26 | 56 | F | L | Right temporal | 107 | 92 | 96 | 109 | 99 | 15* | 6* | 26* | 17* | 35* | 33* | 33 | 16.5 | 12 |
| C27 | 45 | M | R | Left temporal | 79 | 61 | 57 | 80 | 66 | n.a. | n.a. | n.a. | 1* | 17* | 11* | 28 | 4 | 6 |
| C34 | 70 | M | L | Bilateral temporal | n.a. | n.a. | n.a. | n.a. | n.a. | n.a. | n.a. | 13* | 2* | 11* | 0* | 1%ile | 1%ile | 1%ile |
| H42 | 29 | M | R | Not localized | 87 | 75 | 78 | 91 | 79 | 16 | 6 | 22 | 14 | 37 | 36 | 36 | 14.5 | 14.5 |
| H43 | 27 | F | L | Left temporal | n.a. | n.a. | 84 | 86 | n.a. | 19 | 8 | 18 | 17 | 30 | 24 | 23.5 | 14.5 | 14.5 |
| H44 | 58 | F | L | Right temporal | 74 | 77 | 72 | 78 | 74 | 12 | 5 | 10 | 3 | 34 | 28 | 29 | 8 | 7 |
| H45 | 41 | M | R | Not localized | n.a. | n.a. | 114 | 91 | n.a. | 14 | 3 | 23 | 22 | 37 | 30 | 35 | 24.5 | 26 |
| H47 | 20 | M | L | Right amygdala | 105 | 97 | 94 | 111 | 100 | 18 | 8 | 20 | 14 | 36 | 34 | 36 | 23.5 | 23.5 |
| H48 | 54 | M | R | Left temporal | n.a. | n.a. | 76 | 89 | n.a. | 0 | 0 | 13 | 8 | 28 | 22 | 33 | 10.5 | 16.5 |

**(continued)**

| **ID** | **Nr Amygdala Neurons** | | | | | | | | **Nucleus** | |
| --- | --- | --- | --- | --- | --- | --- | --- | --- | --- | --- |
|  | **Total** | **>0.2 Hz** | **Left** | **Right** | **Fear** | **Happy** | **Unam** | **Am** | **Left** | **Right** |
| C26 | 32 | 27 | 27 | 0 | 4 | 0 | 10 | 0 | BL | BL |
|  | 30 | 17 | 17 | 0 | 7 | 0 | 5 | 0 |  |  |
| C27 | 22 | 15 | 15 | 0 | 1 | 3 | 1 | 0 | Ce | BL |
|  | 2 | 2 | 2 | 0 | 0 | 0 | 0 | 0 |  |  |
| C34 | 10 | 9 | 5 | 4 | 1 | 0 | 0 | 0 | BL | BL |
| H42 | 40 | 29 | 29 | 0 | 2 | 3 | 1 | 0 | n.a. | n.a. |
| H43 | 21 | 17 | 17 | 0 | 1 | 1 | 1 | 0 | BL | La |
|  | 12 | 8 | 8 | 0 | 0 | 3 | 0 | 0 |  |  |
|  | 23 | 8 | 8 | 0 | 2 | 0 | 0 | 0 |  |  |
| H44 | 6 | 6 | 0 | 6 | 0 | 0 | 0 | 2 | n.a. | n.a. |
| H45 | 6 | 6 | 0 | 6 | 0 | 0 | 2 | 0 | n.a. | n.a. |
| H47 | 41 | 29 | 14 | 15 | 0 | 0 | 5 | 0 | BL | BL |
| H48 | 32 | 22 | 5 | 17 | 1 | 2 | 1 | 0 | BL | BL |
|  | 50 | 39 | 18 | 21 | 2 | 0 | 3 | 1 |  |  |

Abbreviations: Hand: Dominant handedness; WAIS-III: IQ scores from the Wechsler Adult Intelligence Scale: performance IQ (PIQ), verbal IQ (VIQ), full scale IQ (FSIQ), perceptual organization index (POI), verbal comprehension index (VCI). All WAIS-III scores are on average 100 with a standard deviation of 15 in the normal population (69 and less falls in the clinically abnormal range, 70-79 borderline, 80-89 low average, 90-109 average, 110-119 high average, 120-129 superior, and 130+ very superior). WMS-R and WMS-III are the Wechsler memory scale revised and version 3, respectively. Subtests are verbal paired associates (VPA), logical memory (LM) and visual reproduction (Vis). 1 and 2 are immediate and delayed, respectively. Scores are raw scores. Scores from the Rey-Osterrieth Complex Figure test are raw scores (except C34 as percentiles) from the subtests copy (visuospatial perception and construction), immediate recall reproduction (IR, additional short-term visual memory demands), and 30-minute delayed recall reproduction (DR, additional longer-term visual memory demands). Tests indicated with n.a. were not performed for clinical reasons.

Patients C26, C27 and H48 performed two sessions and patient H43 performed three sessions (Each row of neurons represent a separate recording session. Each session was recorded on a separate day). >0.2 Hz: neurons that had an overall firing rate greater than 0.2Hz. These neurons were included for further analysis. Left: neurons that were recorded from the left amygdala and had a firing rate greater than 0.2Hz. Right: neurons that were recorded from the right amygdala and had a firing rate greater than 0.2Hz. Fear: emotion-tracking neurons that had higher firing rate for more fearful faces. Happy: emotion-tracking neurons that had higher firing rate for happier faces. Unam: ambiguity-coding neurons that had higher firing rate for unambiguous faces. Am: ambiguity-coding neurons that had higher firing rate for more ambiguous faces. BL: basolateral amygdaloid nucleus. Ce: central amygdaloid nucleus. La: lateral amygdaloid nucleus. Nuclei indicated with n.a. were not mapped due to missing T1 structural images (either pre- or post- operative images).

| Experiments | Models | DIC value |
| --- | --- | --- |
| *Experiment 1a: Emotion judgement* | *v ~ 1 + C(Category) : C(Ambiguity) (Categorical ambiguity)* | 4867 |
| *Experiment 1b: Gender judgement* | *v ~ 1 + C(Category) : C(Ambiguity) (Categorical ambiguity)* | 5485 |
| *Experiment 1c: Wealth judgement* | *v ~ 1 + C(Category) : C(Ambiguity) (Categorical ambiguity)* | 5724 |

**Table S2. Behavioral models with separate stimulus categories and objective ambiguity levels.**

**Table S3. Comparisons between models with threshold modulation and models with drift rate modulation only.**

| Behavioral models with varying boundary separation (threshold). | | DIC value |
| --- | --- | --- |
| Behavioral  (Experiment 1a) | *Threshold (a) ~ 1+ Ambiguity (Model 13)* | 6181 |
|  | *Threshold (a) ~ 1+ Category + Ambiguity (Model 14)* | 6186 |
|  | *Threshold (a) ~ 1+ Category : Ambiguity (Model 15)* | 6185 |
|  | *Threshold (a) ~ 1+ Category + Ambiguity + Category : Ambiguity (Model 16)* | 6186 |
| EEG models with varying boundary separation. | | |
| EEG  (Experiment 1a) | *Threshold (a) ~ 1+ LPP (Model 17)* | 6219 |
|  | *Threshold (a) ~ 1+ Category + LPP (Model 18)* | 6217 |
|  | *Threshold (a) ~ 1+ Category : LPP (Model 19)* | 6218 |
|  | *Threshold (a) ~ 1+ Category + LPP + Category : LPP (Model 20)* | 6219 |
| Behavioral models with both drift rates and thresholds. | | |
| Behavioral  (Experiment 1a) | *v ~ 1+ Category + Ambiguity + Category : Ambiguity, a ~ 1+ Ambiguity (Model 21)* | 4695 |
|  | *v ~ 1+ Category + Ambiguity + Category : Ambiguity (Model 4)*  ***(Best model)*** | 4693 |
| EEG models with both the drift rate and threshold. | | |
| EEG  (Experiment 1a) | *v ~ 1+ Category + LPP + Category : LPP, a ~ 1+ LPP (Model 22)* | 4863 |
|  | *v ~ 1+ Category + LPP + Category : LPP (Model 8)* ***(Best model)*** | 4867 |

### Supplementary Figures

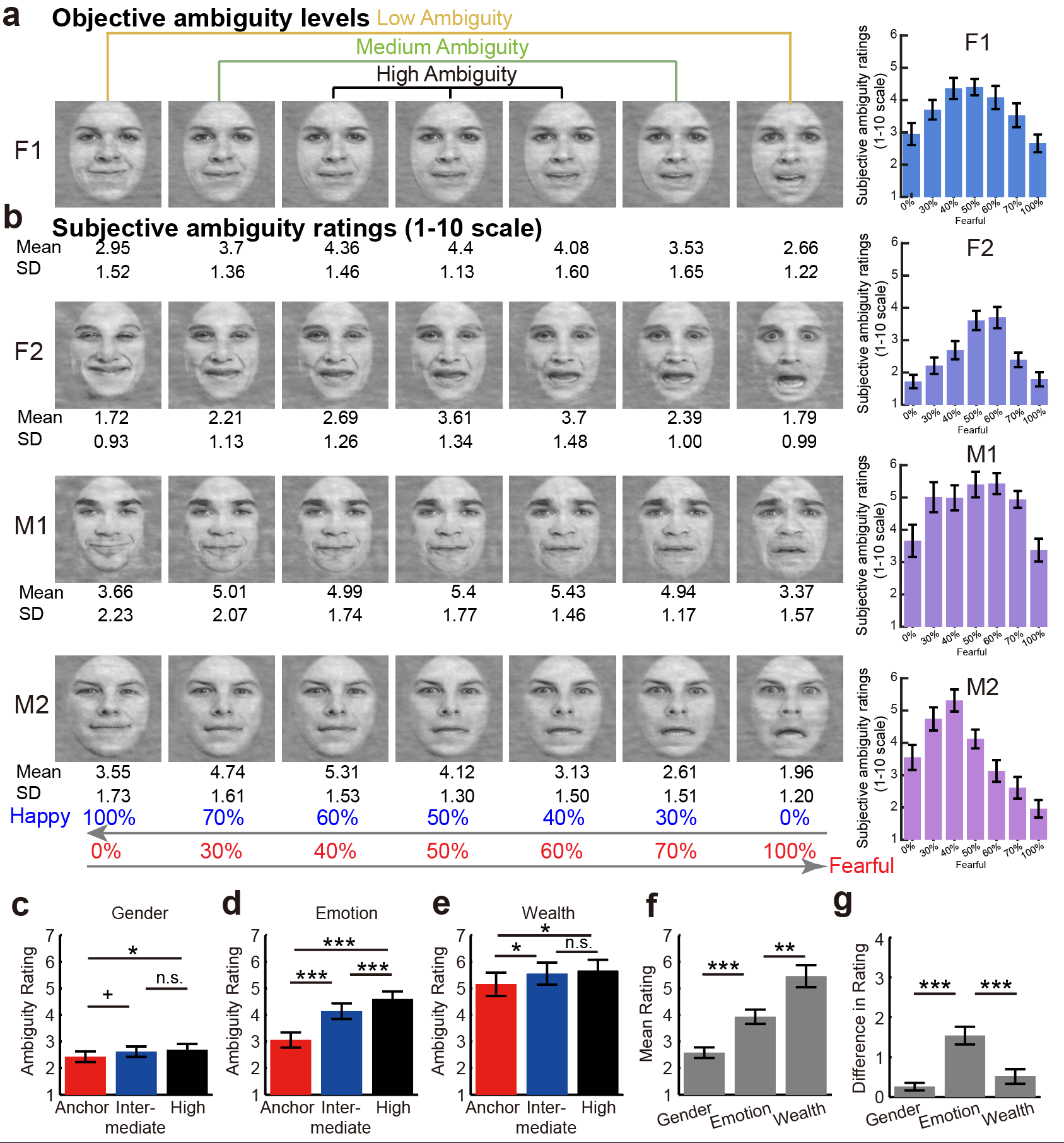

**Figure S1. Subjective ambiguity ratings for each stimulus, across levels and tasks.** (**a**) Morphed emotional face stimuli (F1, F2, M1, M2) vary from 100% happy to 100% fearful. (**b**) Subjective ambiguity ratings (1–10 scale) show a consistent inverted-U pattern across identities, with the highest perceived ambiguity at intermediate morph levels. Error bars indicate ± s.e.m. **(c–e) Ambiguity ratings are shown for Gender, Emotion, and Wealth judgments across anchor, intermediate, and high ambiguity levels. The Wealth task received the highest overall ambiguity ratings (e, f), whereas the Gender task showed the lowest (c, f). The Emotion task exhibited the largest separation across ambiguity levels (d), consistent with the direct stimulus–response mapping that makes ambiguity manipulation most perceptually salient. Difference scores (g) confirm that ambiguity effects were strongest for Emotion, moderate for Wealth, and minimal for Gender. Error bars indicate ± s.e.m.** *p < .05, ** p < .01**, *** *p* < .001**, ^+^*p* < .10, n.s. indicates not significant. Adapted from Wang et al., 2017, Nature Communications.

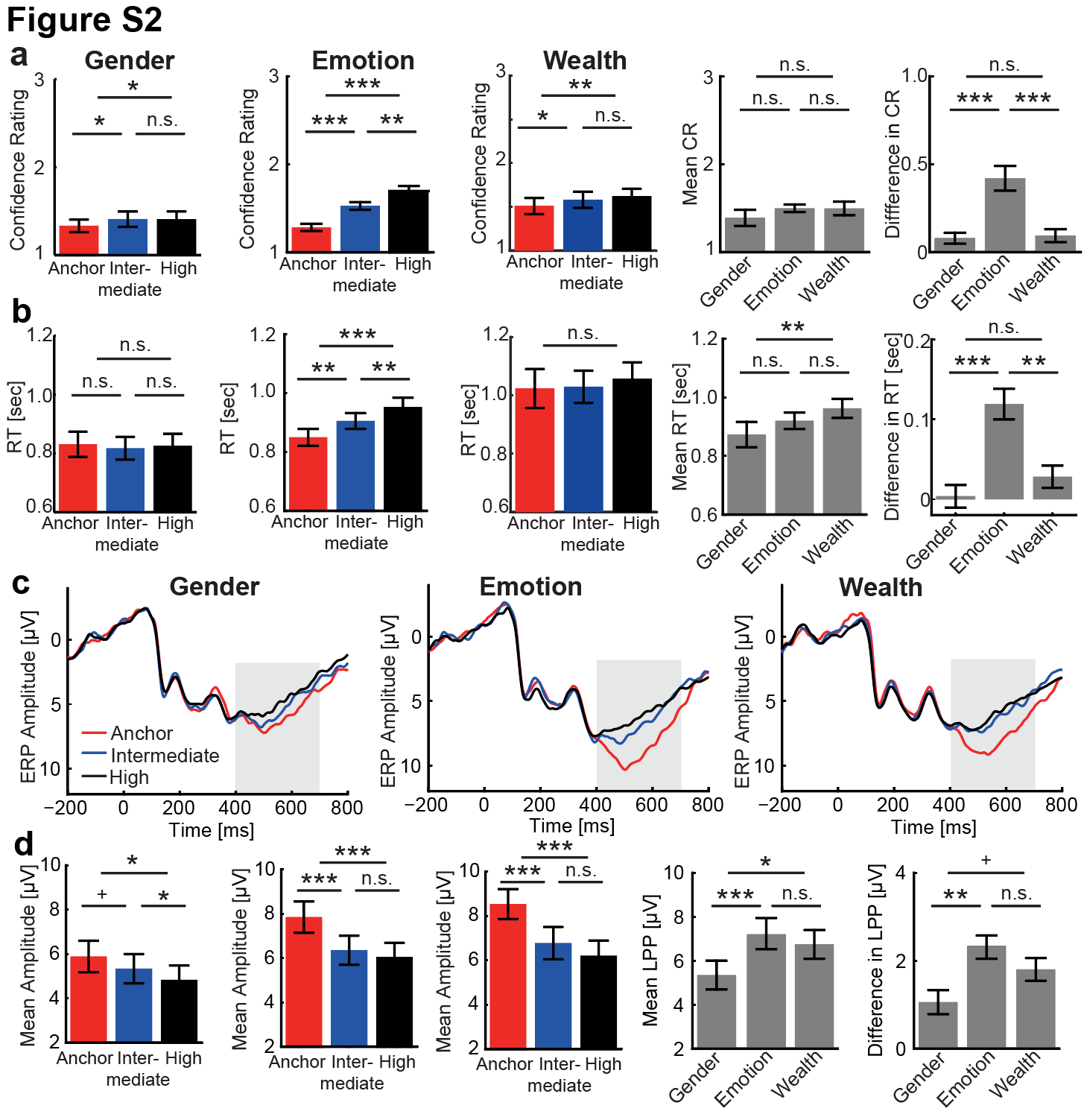

**Figure S2. Behavioral and ERP effects across Gender, Emotion, and Wealth judgments in Experiment 1**. **(a-b)** Confidence rating (CR, where 1 = most confident, 3 = least confident) and reaction time (RT) vary systematically with ambiguity level (Anchor = no ambiguity, Intermediate = moderate ambiguity, High = highest ambiguity). The strongest effects of ambiguity on confidence ratings and reaction times (high > anchor) were observed in the facial expression judgment task, where the ambiguity was relevant to the task. (**c**) ERP waveforms at Pz show corresponding modulation in the late positive potential (LPP; 400–700 ms window, shaded). (d) Bar plots summarize mean amplitudes and LPP responses, demonstrating robust ambiguity-related effects for Emotion and Wealth, and modest effect for Gender. Cross-task comparisons reflect the mean and difference in CR, RT, and LPP amplitudes across ambiguity levels. Adapted from Sun et al., 2017, NeuroImage; Sun et al., 2017, eNeuro.

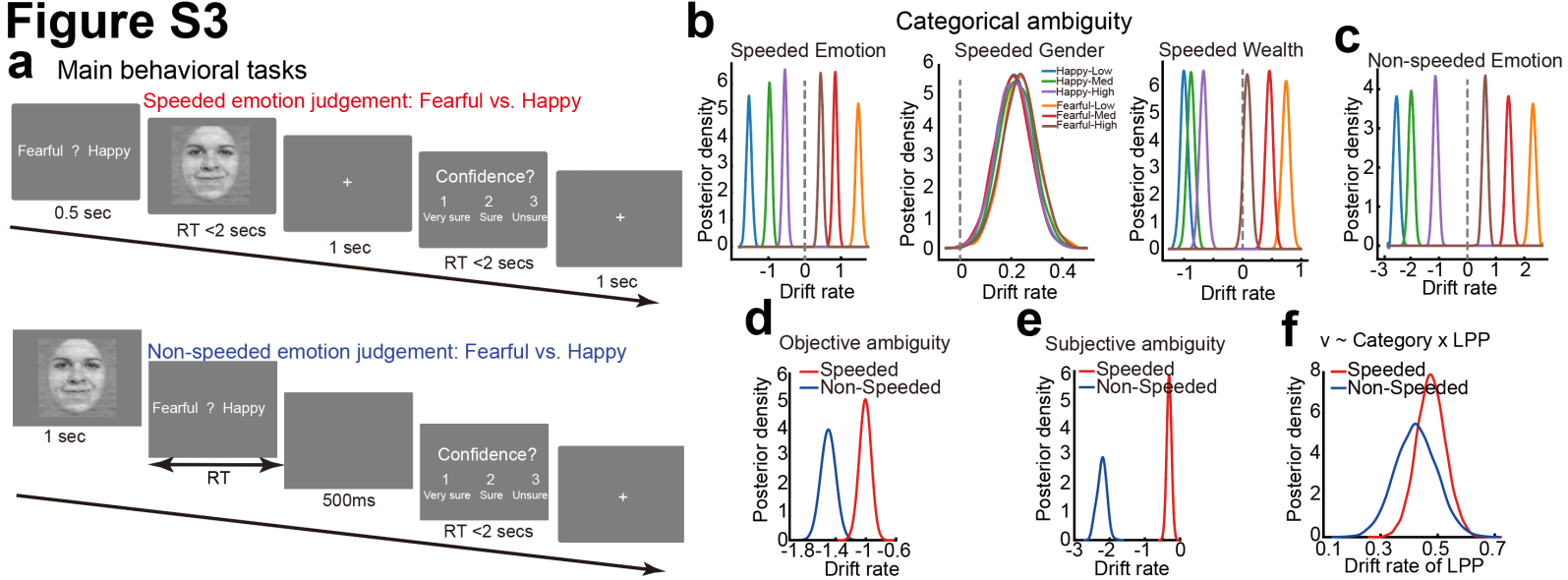

**Figure S3. Additional task procedure and modeling results corresponding to Figure 2.** (**a**) Comparison of tasks with speeded responses (Experiment 1a) and non-speeded responses (Experiment 2a). Note that the facial images shown here were adapted from a publicly available database. (**b**) Posterior drift-rate distributions by emotion category and ambiguity level across the emotion, gender, and wealth judgment tasks. Extension on Figure 2. Happy (blue, green, purple) distributions cluster on the negative side of the drift axis, whereas fearful (brown, red, orange) distributions lie on the positive side, illustrating that the sign of the interaction term reflects directional coding for both the emotion and wealth tasks. The width and separation of the distributions capture differences in evidence accumulation strength across ambiguity levels—expressed as an interaction effect—with the emotion task showing the largest separation, which becomes weaker in the wealth judgment. In contrast, the gender task shows no discernible differences in evidence accumulation across morphing levels or stimulus categories. (**c**) Posterior distribution of the category-by-objective ambiguity interaction effect for speeded (red, same as the red distribution in Figure. 2b) and non-speeded (blue, same as the red distribution in Figure. 3b) tasks. Here, objective ambiguity levels are researcher-defined. (**d**) Posterior distribution of the category-by-subjective ambiguity interaction effect for speeded (red) and non-speeded (blue) tasks. Here, subjective ambiguity levels were obtained from an independent group of participants. (**e**) The category × LPP interaction effect on drift rate showed similar patterns for the speeded and non-speeded tasks.

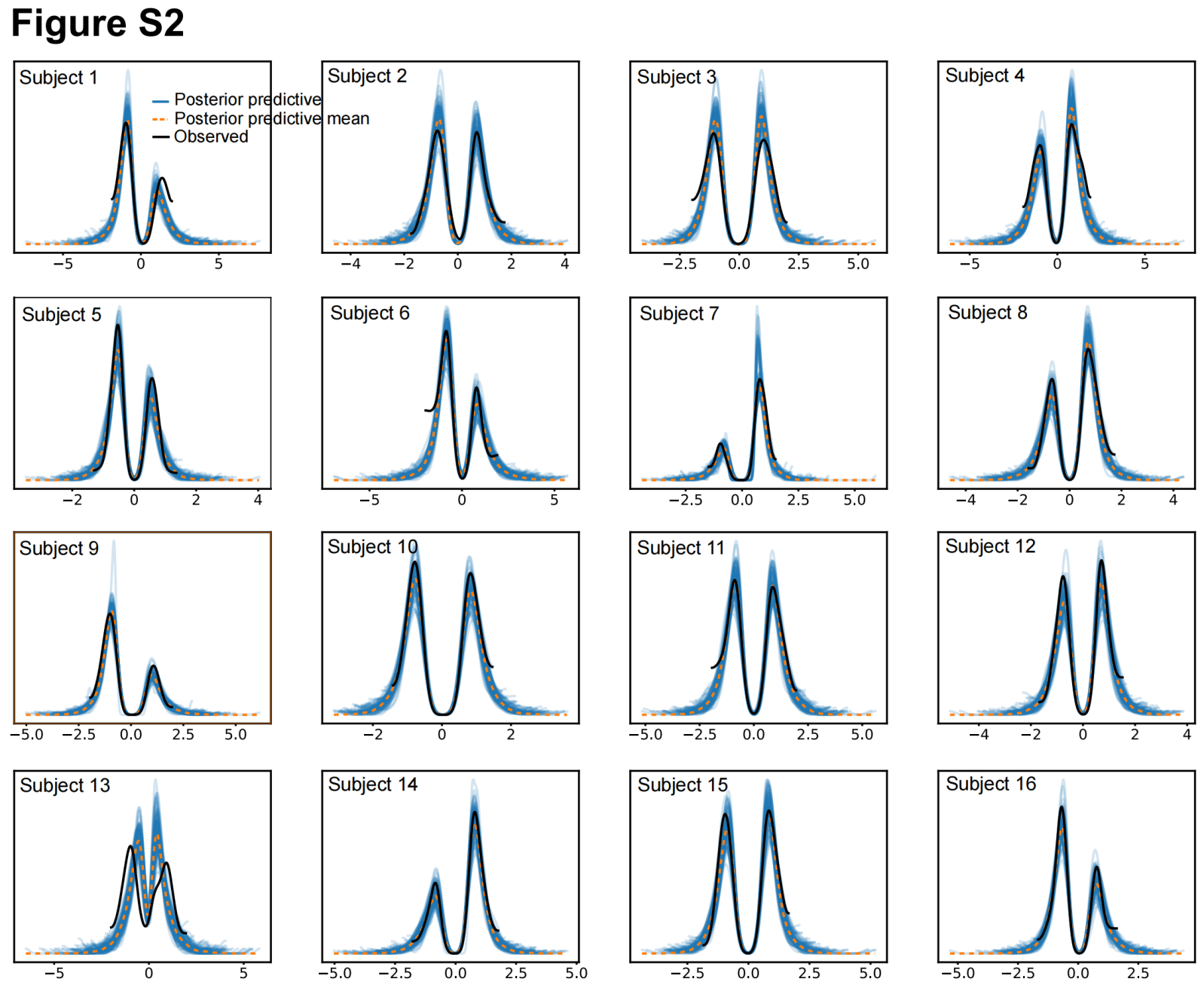

**Figure S4. Posterior predictive reaction times (RTs) and comparison with observed RTs for each subject in the main task (Experiment 1a).** Each panel represents one of the 16 individual subjects in Experiment 1a. The blue lines show posterior predictive RT distributions generated from 100 samples, which are comparable to the observed RT distributions. Overall, the model demonstrates good recovery of RT distributions, with simulated RTs closely aligning with the observed data.

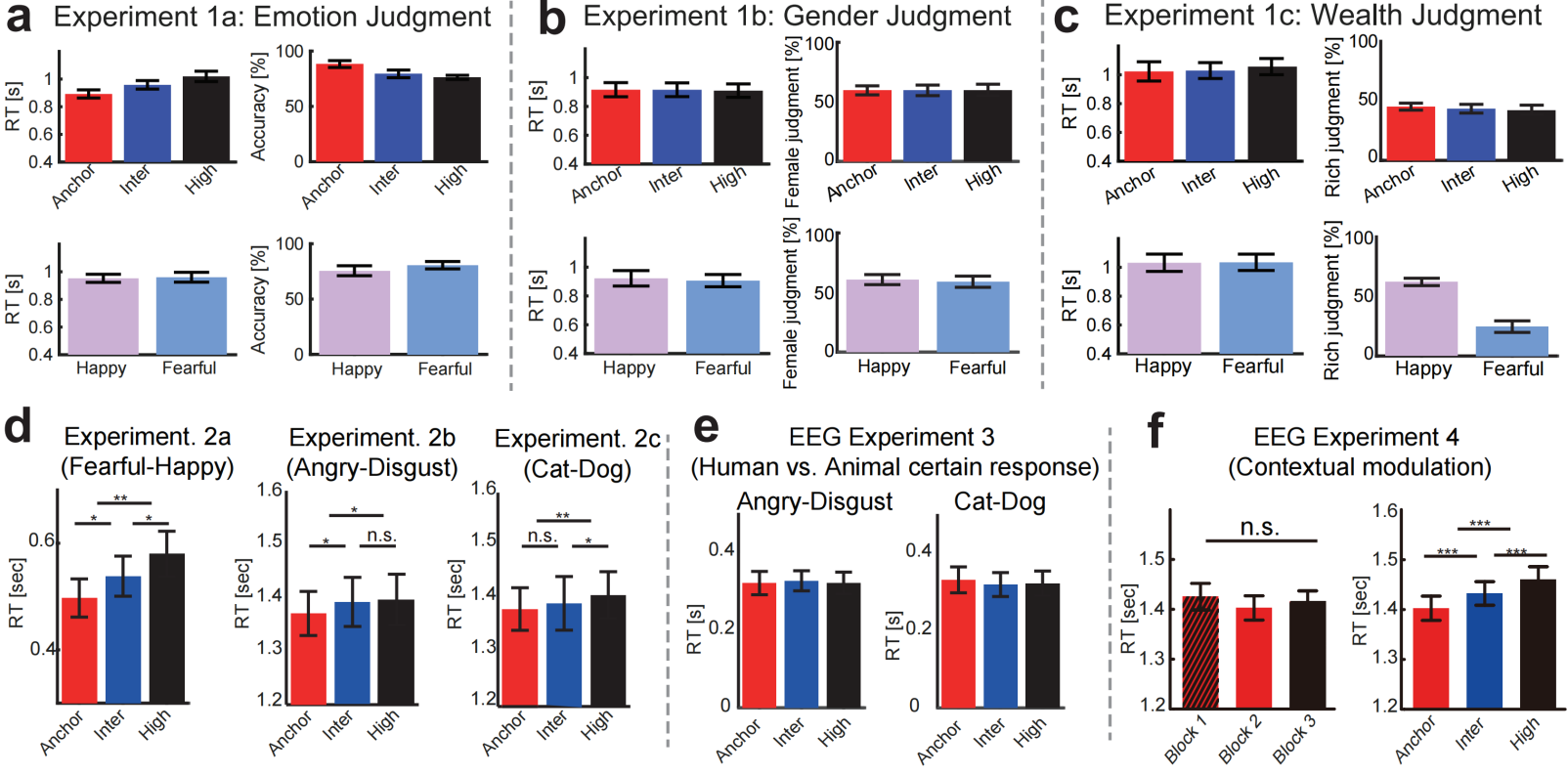

**Figure S5. Behavioral results across Experiments 1-4.** (**a-c**) RTs and behavioral responses across ambiguity levels for Experiment 1 (emotion, gender, and wealth judgments). In the emotion task (**a**), RTs increased and accuracy decreased with higher ambiguity, with no differences between fearful and happy faces. For gender (**b**) and wealth (**c**) judgments, we report the percentage of choosing one category (e.g., “female”, “rich”), which varied with ambiguity and stimulus category; happy faces were more often judged as “rich”. (**d**) In Experiments 2a-c (Fearful-Happy, Angry-Disgust, Cat-Dog), RTs increased reliably with ambiguity, showing robust ambiguity-dependent slowing across tasks with no category-specific differences. (**e**) In Experiment 3 (certain-response EEG task), RTs showed minimal modulation, consistent with the design requiring responses only when judging whether the stimulus was an animal or a human. (**f**) In Experiment 4 (contextual modulation EEG task), RTs did not differ across low-ambiguity blocks but increased systematically with ambiguity in the mixed-ambiguity block, confirming ambiguity-dependent slowing. Error bars indicate ±s.e.m. Adapted from Sun et al., 2017, NeuroImage; Sun et al., 2017, eNeuro.

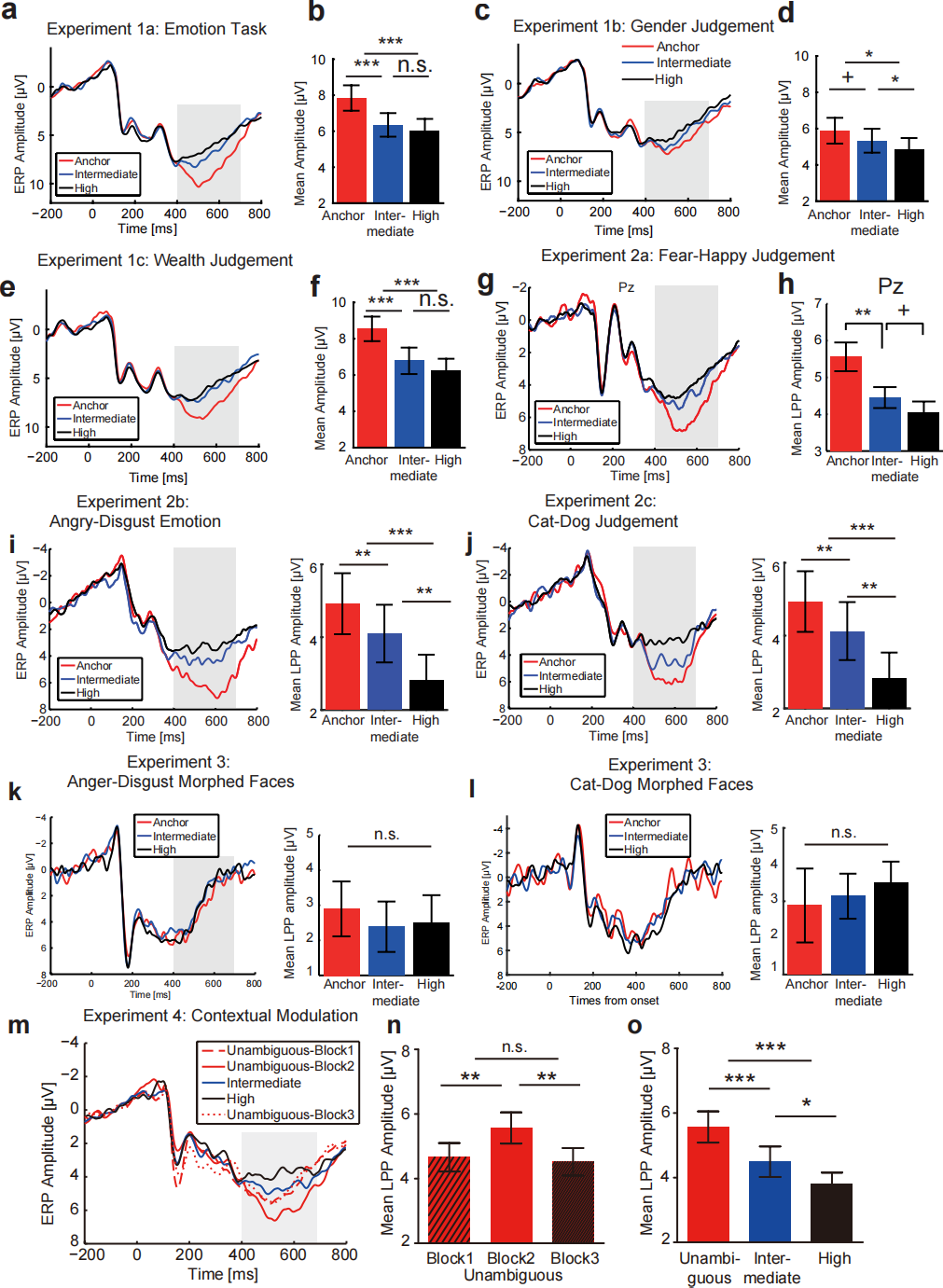

**Figure S6. Grand-average ERPs by ambiguity levels across Experiments 1 to 4.** (**a–b**) In Experiment 1a (Emotion Task), late positive potentials (LPP; 400–700 ms) increased with higher ambiguity relative to low ambiguity levels. (**c–d**) Experiment 1b (Gender Judgment) showed minimal modulation of LPP amplitude by emotional ambiguity. (**e–f**) Experiment 1c (Wealth Judgment) exhibited a similar ambiguity-dependent enhancement of LPP amplitude observed in the emotion task. (**g–h**) In Experiment 2a (Non-speeded Fearful–Happy Judgment), Pz LPP amplitude increased with ambiguity. **(i)** Experiment 2b (Angry–Disgust Judgment) showed clear LPP differences across ambiguity levels. **(j)** Experiment 2c (Cat–Dog Judgement) also showed ambiguity-related LPP modulation. (**k–l**) In Experiment 3 (morphed Angry–Disgust and Cat–Dog stimuli, with a certain response prompt), ambiguity levels did not reliably modulate LPP amplitude. (**m–o**) In Experiment 4 (contextual modulation), ERP waveforms showed no systematic block-wise differences in low ambiguous trials. However, LPP amplitude varied as a function of ambiguity levels. Across experiments, ambiguity-dependent increases in LPP amplitude were consistently observed in tasks involving emotional or socially meaningful categories, but not in decisions with low ambiguity levels. Error bars indicate ± s.e.m.; *p < .05, **p < .01, ***p < .001, n.s. indicates not significant. Adapted from Sun et al., 2017, NeuroImage; Sun et al., 2017, eNeuro.
